# Human Skeletal Muscle Fiber Heterogeneity Beyond Myosin Heavy Chains

**DOI:** 10.1101/2023.09.07.556665

**Authors:** Roger Moreno-Justicia, Thibaux Van der Stede, Ben Stocks, Jenni Laitila, Robert A Seaborne, Alexia Van de Loock, Eline Lievens, Diana Samodova, Leyre Marín-Arraiza, Oksana Dmytriyeva, Robin Browaeys, Kim Van Vossel, Lukas Moesgaard, Nurten Yigit, Jasper Anckaert, Anneleen Weyns, Ruud Van Thienen, Ronni E Sahl, Edmar Zanoteli, Michael W Lawlor, Michael Wierer, Pieter Mestdagh, Jo Vandesompele, Julien Ochala, Morten Hostrup, Wim Derave, Atul S Deshmukh

## Abstract

Skeletal muscle is an inherently heterogenous tissue comprised primarily of myofibers, which are historically classified into three distinct fiber types in humans: one “slow” (type 1) and two “fast” (type 2A and type 2X), delineated by the expression of myosin heavy chain isoforms (MYHs). However, heterogeneity between and within traditional fiber types remains underexplored. Indeed, whether MYHs are the main classifiers of skeletal muscle fibers has not been examined in an unbiased manner. Through the development and application of novel transcriptomic and proteomic workflows, applied to 1050 and 1038 single muscle fibers from human *vastus lateralis*, respectively, we show that MYHs are not the only principal drivers of skeletal muscle fiber heterogeneity. Instead, metabolic, ribosomal, and cell junction proteins are a source of multi-dimensional variation between skeletal muscle fibers. Furthermore, whilst slow and fast fiber clusters can be identified, described by their contractile and metabolic profiles, our data suggests that type 2X fibers are not phenotypically distinct to other fast fibers at an omics level. Moreover, MYH-based classifications do not adequately describe the phenotype of skeletal muscle fibers in one of the most common genetic muscle diseases, nemaline myopathy, with fibers shifting towards a non-oxidative phenotype independently of MYH-based fiber type. We also characterize novel transcriptomic and proteomic features of slow and fast skeletal muscle fibers, including identifying several muscle fiber type-specific polypeptides, termed microproteins, encoded by transcripts annotated as non-coding RNA. Overall, our data indicates that skeletal muscle fiber heterogeneity is multi-dimensional with sources of variation beyond myosin heavy chain isoforms.

## Introduction

Cellular heterogeneity is an inherent feature of all biological systems, allowing for cellular specialization to meet the diverse demands imposed upon tissues and cells^1^. The classic view of skeletal muscle fiber heterogeneity is that motoneurons define the typology of the fibers within a motor unit, with fiber types (i.e. type 1, type 2A, and type 2X in humans) being defined by myosin heavy chain isoform (MYH) characteristics^2^. This was first based on its ATPase pH lability^3,4^ and later by its molecular MYH expression^5^. However, skeletal muscle fibers are increasingly being viewed along a continuum, as opposed to discrete fiber types^6^, fueled by the identification and subsequent acceptance of the existence of “hybrid” fibers that express varying proportions of multiple MYHs simultaneously. Nonetheless, the field still relies heavily on MYHs as primary classifiers of muscle fibers, a perspective that might be limited and heavily biased by early studies in rodents, displaying a different MYH expression profile and thus fiber type range than humans^2^. The picture is further complicated by different skeletal muscles displaying specialized fiber type profiles within humans^7^. The *vastus lateralis* is a mixed muscle type with average (and therefore representative) MYH expression profiles^7^. Together with this, its accessibility for sampling makes it the most commonly studied muscle in humans.

An unbiased exploration of skeletal muscle fiber diversity with powerful omics tools is thus heavily warranted, but challenging, in part owing to the multinucleated nature of skeletal muscle fibers. Nonetheless, both transcriptomics^8,9^ and proteomics^10^ technologies have recently experienced a sensitivity revolution due to various technological advances, making profiling of skeletal muscle at a single fiber resolution now possible. As a result, notable progress has already been made in describing the single muscle fiber diversity and responses to atrophic stimuli and aging^11–18^. Importantly, these technological advances lend themselves to being advantageous in the clinical setting, helping to describe in greater detail and precision the dysregulation associated with disease. For example, the underlying pathophysiology of nemaline myopathy, one of the most prevalent genetic muscle disorders (MIM 605355 and MIM 161800), is complex and convoluted^19,20^, thus a deeper characterization of skeletal muscle fiber dysregulation may spur significant developments in our understanding of the disease.

Our methodological development of transcriptome and proteome profiling of single skeletal muscle fibers manually isolated from human biopsy specimens, and their application to over a thousand fibers each, enabled us to investigate the cellular heterogeneity of human skeletal muscle fibers. In doing so, we demonstrate the power of muscle fiber phenotyping at the transcriptomic and proteomic level, identifying that metabolic, ribosomal, and cell junction proteins are important sources of variation among muscle fibers. Furthermore, with our proteomics workflow we characterize the clinical implications of nemaline myopathies within single skeletal muscle fibers, revealing a coordinated shift towards non-oxidative fibers independently of MYH-based fiber type.

## Results

### Development of high-sensitivity and high-throughput single muscle fiber transcriptomic and proteomic pipelines

To investigate the heterogeneity of human skeletal muscle fibers, we developed two novel workflows to enable transcriptome and proteome profiling of single skeletal muscle fibers (**Fig 1A and 1 S1A**). Several methodological steps were developed and optimized, from sample storage and preservation of RNA and protein integrity to optimizing the throughput of each method. This was achieved for transcriptome analysis by inserting sample-specific molecular barcodes during the initial reverse transcription step, enabling pooling of 96 fibers for further efficient downstream processing. Rich transcriptome data was further obtained by deeper sequencing (± 1M reads per fiber) compared to conventional single cell methods^21^. For proteomics, the combination of a short chromatographic gradient (21 minutes) and timsTOF DIA-PASEF-based mass spectrometry was used to optimize proteome depth, whilst maintaining a high-throughput^22,23^. To investigate skeletal muscle fiber heterogeneity in the healthy state, the transcriptome was determined for 1050 individual fibers from 14 healthy adult donors, whilst the proteome was determined for 1038 fibers from 5 healthy adult donors (**Table S1)**. These datasets will be referred to as the 1000 fiber transcriptome and proteome datasets, respectively, throughout this manuscript. Our approach detected a total of 27237 transcripts and 2983 proteins in the 1000 fiber transcriptomics and proteomics studies (**Fig 1A, Table S2-3**). After filtering for >1000 detected genes and for 50% valid values within each fiber in both transcriptomics and proteomics datasets, respectively, downstream bioinformatic analyses were performed on 925 and 974 fibers in the transcriptome and proteome, respectively. On average 4257 ± 1557 genes and 2015 ± 234 proteins (mean ± SD) were detected per fiber after filtering, with limited inter-individual variation (**Fig 1 S1B-C, Table S4-5**). The intra-individual variation within a participant was more substantial however, most likely due to differences in RNA/protein yield among fibers of different length and cross-sectional area. For the majority of proteins (>2000), the coefficient of variation was below 20% (**Fig 1 S1D**). Both methodologies captured a wide dynamic range of transcripts and proteins, with features known to be important for muscle contraction being highly expressed (e.g. ACTA1, MYH2, MYH7, TNNT1, TNNT3) (**Fig 1 S1E-F**). A large part of the detected features were shared between the transcriptome and proteome datasets (**Fig 1 S1G**), alongside a reasonable correlation (r = 0.52) in average UMI counts/LFQ intensities for these features (**Fig 1 S1H**).

**Figure 1.**
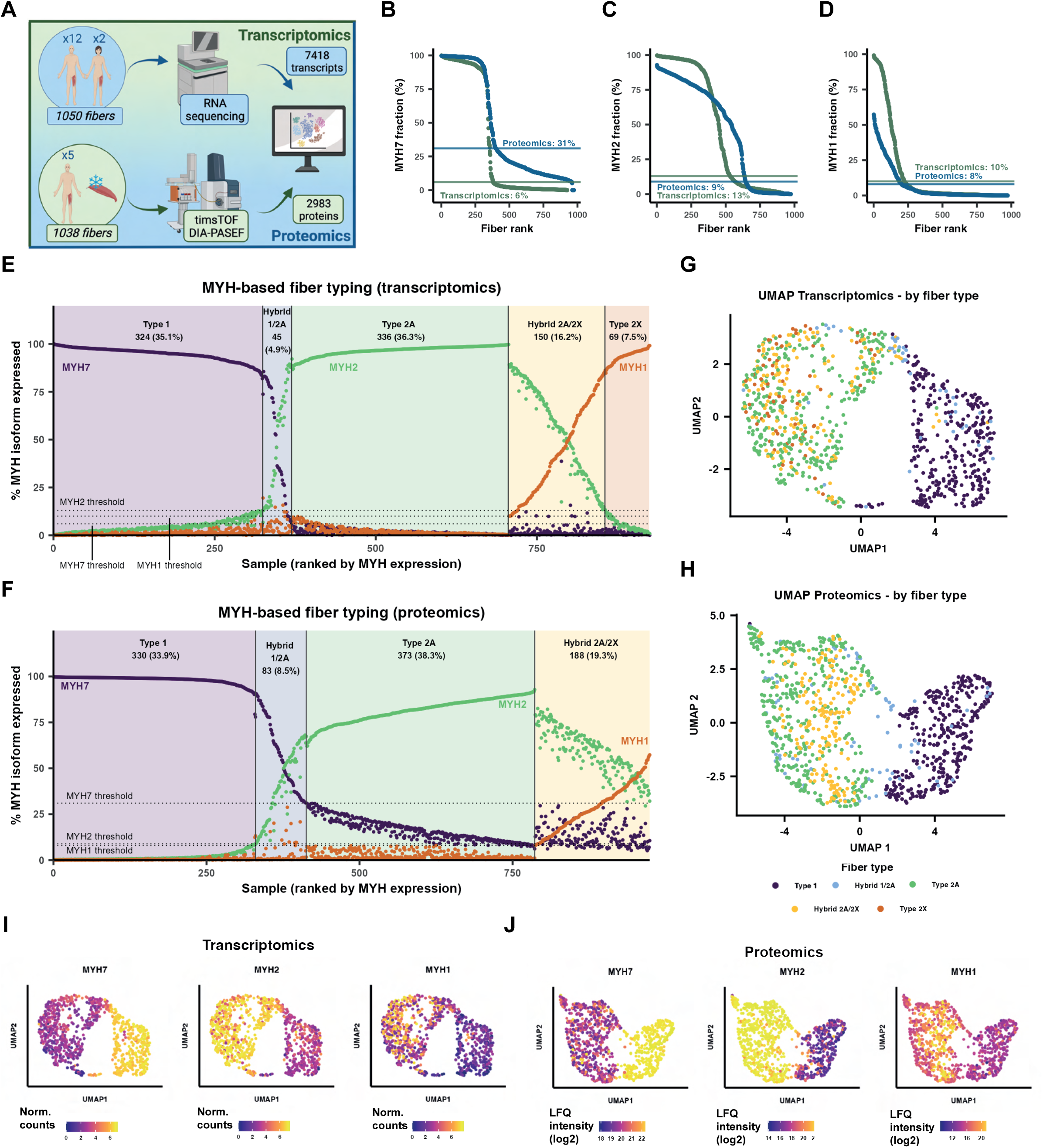
Transcriptomic- and proteomic-based clustering reveals only two distinct fiber types. **(A)** Transcriptomic and proteomic workflow. **(B-D)** Dynamic range curves for MYH7, MYH2, and MYH1, with calculated assignment thresholds for fiber typing. **(E-F)** Distribution of fibers by expression of MYHs in transcriptomics and proteomics datasets. **(G-H)** Uniform manifold approximation and projection (UMAP) plots for transcriptomics and proteomics, colored by MYH-based fiber type. **(I-J)** Feature plots displaying the expression of MYH7, MYH2, and MYH1 in transcriptomics and proteomics.

### Type 2X is not a distinct fiber type

We initially set out to define the MYH-based fiber type of each fiber using an optimized methodology, leveraging the high sensitivity and dynamic range of MYH expression in the omics datasets. Previous studies have used arbitrary cut-offs to assign a fiber as pure type 1, type 2A, type 2X, or hybrid, based on a fixed percentage of expression for the different MYHs^11,14,24^. We employed a different approach, in which we ranked the expression of each fiber by the MYHs used for fiber typing: MYH7, MYH2 and MYH1, corresponding to type 1, type 2A and type 2X fibers, respectively. We then mathematically calculated the bottom knee for each of the resulting curves and used it as a threshold to assign a fiber as being positive (above threshold) or negative (below threshold) for each MYH (**Fig 1B-D**). These data show that MYH7 (**Fig 1B**) and MYH2 (**Fig 1C**) have a more pronounced on/off expression profile on the RNA level, compared to the protein level. Indeed, at the protein level, very few fibers did not express MYH7 and no fibers had 100% MYH2 expression. Next, we used the determined expression thresholds to assign MYH-based fiber types for all fibers in each dataset. For example, a MYH7^+^/MYH2^-^/MYH1^-^ was assigned as type 1, and a MYH7^-^/MYH2^+^/MYH1^+^ was assigned as a hybrid type 2A/2X fiber (see **Table S6** for full description). When combining all fibers, a very similar MYH-based fiber type distribution was observed at the RNA (**Fig 1E**) and protein (**Fig 1F**) levels, with an expected inter-individual variation in relative MYH-based fiber type composition (**Fig 1 S2A**). Most fibers were considered pure type 1 (34-35%) or type 2A (36-38%), although a substantial number of hybrid 2A/2X fibers (16-19%) were also detected. A striking discrepancy was that pure type 2X fibers could only be detected at the RNA but not at the protein level, indicating that fast MYH expression is, at least in part, post-transcriptionally regulated.

We validated the MYH-based fiber typing from the proteomics data against an antibody-based dot blot technique, in which both methodologies were in 100% agreement on the identification of both pure type 1 and type 2A fibers (**Fig 1 S2B**). However, the increased sensitivity of the proteomics-based approach was superior in identifying hybrid fibers and in quantifying the proportion of each MYH within each fiber. These data demonstrate the efficacy of using an unbiased high-sensitivity omics-based approach for the characterization of skeletal muscle fiber types.

We then utilized the full depth of information that transcriptomics and proteomics provide to classify fibers in an unbiased manner based on their whole transcriptome or proteome. Using uniform manifold approximation and projection (UMAP) for dimension reduction of 6 principal components (**Fig 1 S3 A-B**), we were able to visualize the variation in fibers in the transcriptome (**Fig 1G**) and proteome (**Fig 1H**). Interestingly, fibers did not cluster by participant in either the transcriptomics or proteomics datasets (**Fig 1 S3C-D**), nor by test day (**Fig 1 S3E**), indicating that intra-individual variance in skeletal muscle fibers outweighs inter-individual variance. Two distinct clusters were apparent in the UMAP plots, which represented “fast” and “slow” fibers (**Fig 1G-H**). MYH7^+^ (slow) fibers clustered to the positive side of UMAP1, and MYH2^+^ and MYH1^+^ (fast) fibers clustered to the negative side of UMAP1 (**Fig 1I-J**). No distinction between the various fast MYH-based fiber types (i.e. type 2A, type 2X, or hybrid 2A/2X) could be identified, however, suggesting that when the whole transcriptome or proteome is taken into account, the expression of MYH1 (**Fig 1 I-J**), or other classical markers of type 2X fibers like ACTN3 or MYLK2 (**Fig 1 S4A-B**), does not discriminate between distinct fiber types. Furthermore, in contrast to MYH2 and MYH7, few transcripts or proteins positively correlate with MYH1 (**Fig 1 S4C-H**), suggesting that MYH1 abundance does not adequately reflect the muscle fiber transcriptome/proteome. Similar conclusions can be drawn when assessing the blended expression of the three MYH isoforms on a UMAP level (**Fig 1 S4I-J**). Thus, whilst type 2X fibers can be identified at the transcriptional level based solely on the quantification of MYHs, MYH1^+^ fibers are not distinct from other fast fibers when the whole transcriptome or proteome is considered.

### Considerable skeletal muscle fiber heterogeneity beyond myosin heavy chain isoforms

As an initial exploration of fiber heterogeneity beyond MYHs, we assessed four established slow fiber type-specific proteins: TPM3, TNNT1, MYL3, and ATP2A2^2^. In both transcriptomics (**Fig 2 S1A**) and proteomics (**Fig 2 S1B**) approaches, the slow isoforms exhibited a high, although not perfect, Pearson correlation coefficient with MYH7. Approximately 25 and 33% of the fibers in the transcriptomics (**Fig 2 S1C**) and proteomics (**Fig 2 S1D**) approaches, respectively, were not classified as pure slow fibers by all gene/protein isoforms. Thus, fiber typing based on multiple gene/protein isoforms introduces additional complexity, even with well-known proteins that are assumed to be fiber type-specific. This suggests that fiber typing based on isoforms of a single family of genes/proteins is likely inadequate to capture the true heterogeneity of skeletal muscle fibers.

To further investigate the omics-wide phenotypical variability between human skeletal muscle fibers, we applied unbiased dimensionality reduction by principal component analysis (PCA) to our data (**Fig 2A**). Similarly to the UMAP plot, neither participant nor test day influenced the clustering of fibers at the PCA level (**Fig 2 S2A-C**). MYH-based fiber type was explained by PC2 in both datasets, displaying clusters for slow type 1 fibers and a second cluster containing the fast type 2A, type 2X and hybrid type 2A/2X fibers (**Fig 2A**). These two clusters were bridged in both datasets by a small number of hybrid type 1/2A fibers. As expected, over-representation analysis of the top PC drivers confirmed that PC2 is driven by contractile and metabolic features (**Fig 2B & 2 S2D-E, Table S7-9**). In general, the MYH-based fiber types adequately explain the continual variance along PC2, except for the so-called type 2X fibers, which were spread across the entirety of the transcriptomic fast cluster.

**Figure 2.**
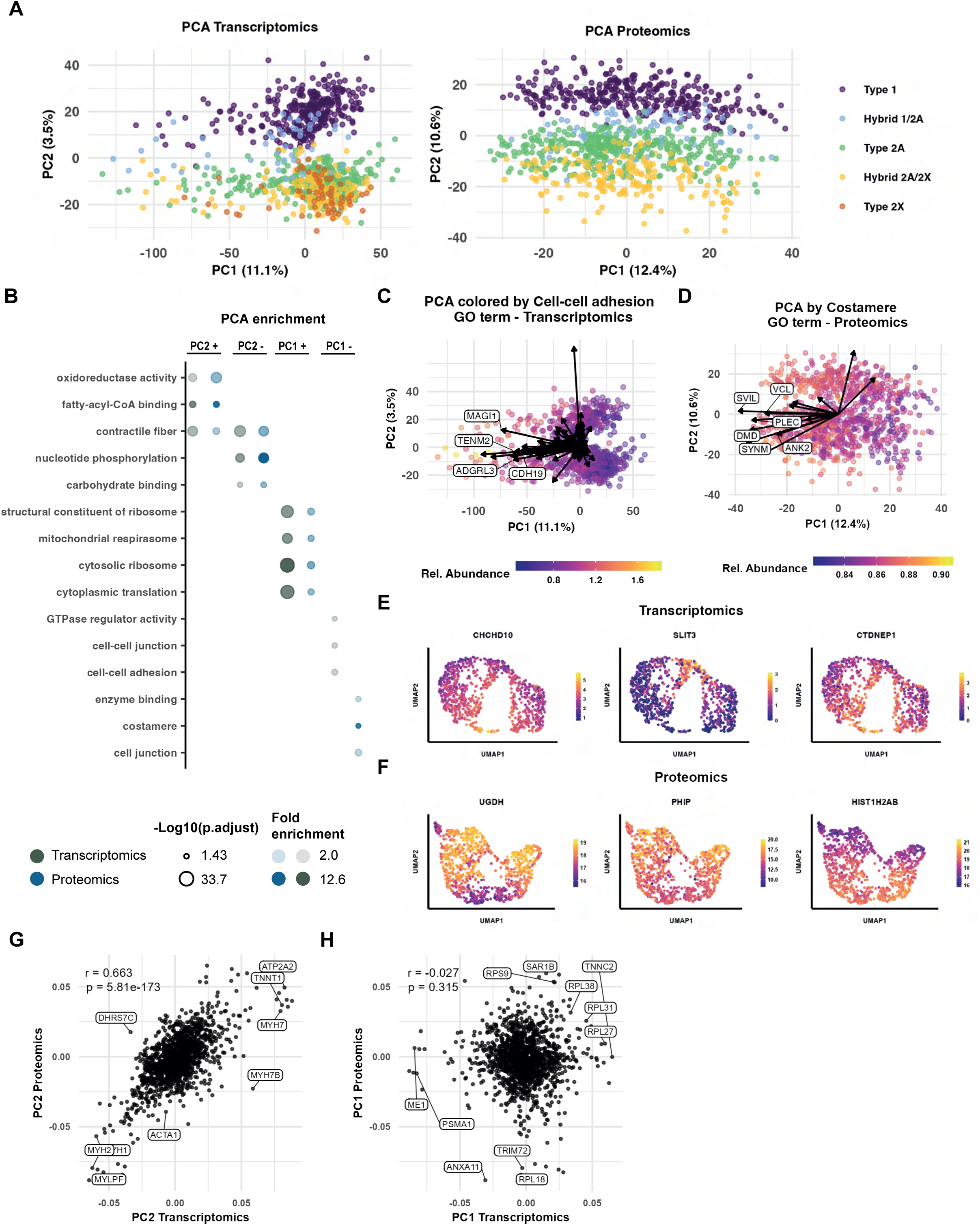
Skeletal muscle fiber heterogeneity beyond MYHs. **(A)** Principal component analysis (PCA) plots of transcriptomics and proteomics datasets colored by MYH-based fiber type. **(B)** Enrichment analysis of transcript and protein drivers of PC2 and PC1. **(C-D)** PCA plot colored by cell-cell adhesion gene ontology (GO) term in the transcriptome, and by costamere GO term in the proteome. Arrows represent transcript and protein drivers and their direction. **(E-F)** Uniform manifold approximation and projection (UMAP) feature plots of clinically relevant features displaying a slow/fast fiber type-independent gradient of expression. **(G-H)** Correlation between drivers of PC2 and PC1 between the transcriptome and proteome.

Unexpectedly, MYH-based fiber type explained only the second greatest degree of variability (PC2), indicating that other biological factors (PC1) independent of MYH-based fiber type, have a substantial role in regulating skeletal muscle fiber heterogeneity. Over-representation analysis of the top drivers in PC1 indicated that variance within PC1 was determined primarily by cell-cell adhesion and ribosomal content in the transcriptome, and costamere and ribosomal proteins in the proteome (**Fig 2B & 2 S2D-E, Table S9**). In skeletal muscle, the costamere connects the Z-disk to the sarcolemma and participates in force transmission and signaling^25^. Annotating PCA plots with cell-cell adhesion (transcriptome, **Fig 2C**) and costamere (proteome, **Fig 2D**) features showed a strong shift to the left side of PC1, suggesting an enrichment for these features in some fibers.

Closer inspection of fiber clustering at the UMAP level showed a MYH-based fiber type-independent gradient of expression for most features, rather than distinct subclusters of muscle fibers. This continuum holds true for several genes related to pathological conditions (**Fig 2E**), such as *CHCHD10* (neuromuscular disorders)^26^, *SLIT3* (muscle loss)^27^ and *CTDNEP1* (muscle disease)^28^. This same continuum was observed in the proteome, with proteins related to neurological diseases (UGDH)^29^, insulin signaling (PHIP)^30^ and transcription (HIST1H2AB) (**Fig 2F**). These data collectively show that slow/fast fiber type independent heterogeneity across fibers is of a continual nature.

### Fiber heterogeneity is post-transcriptionally regulated by ribosomal heterogeneity

Interestingly, drivers of PC2 showed a good correlation (r = 0.663) between the transcriptome and proteome (**Fig 2G**), indicating that the slow and fast fiber types, particularly the contractile and the metabolic profile of skeletal muscle fibers, are transcriptionally regulated. However, there was no correlation between the drivers of PC1 in the transcriptome and the proteome (r = -0.027) (**Fig 2H**). This suggests that slow/fast fiber type-independent variance is for a large part post-transcriptionally regulated. Since PC1 variance was largely explained by ribosomal gene ontology terms and given that ribosomes play a profound and specialized role in the cell by actively participating and influencing protein translation^31^, we next set out to explore this unexpected ribosomal heterogeneity.

We first colored the proteomics PCA plot based on the relative abundance of the proteins within the “cytosolic ribosome” GOCC term (**Fig 3A**). Although the term was enriched on the positive side of PC1, and a slight gradient could be observed accordingly, ribosomal proteins were driving separation in both directions of PC1 (**Fig 3A**). Amongst the ribosomal proteins enriched in the negative direction of PC1 were RPL18, RPS18 and RPS13 (**Fig 3B**), whilst RPL31, RPL35 and RPL38 (**Fig 3C**) were major drivers in the positive direction of PC1. Interestingly, RPL38 and RPS13 are highly expressed in skeletal muscle when compared to other tissues (**Fig 3 S1A**). These distinct ribosomal signatures across PC1 could not be observed in the transcriptome (**Fig 3 S1B**), indicating that this phenomenon is post-transcriptionally regulated.

**Figure 3.**
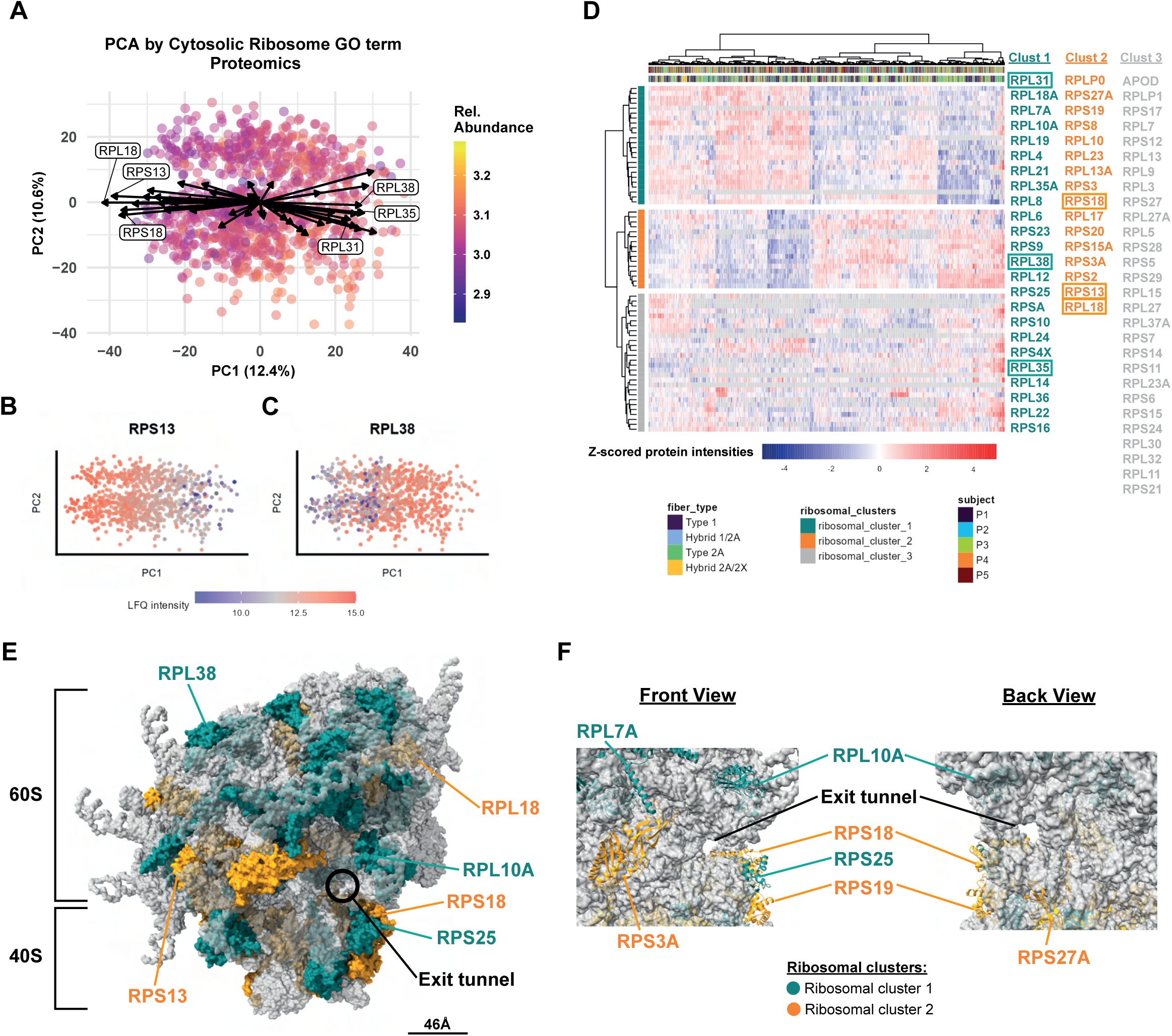
Ribosomal heterogeneity drives fiber type-independent heterogeneity. **(A)** Principal component analysis (PCA) plot colored by cytosolic ribosome gene ontology (GO) term in the proteome. Arrows indicate the direction towards which the proteins are driving the variance in the PCA plot. The length of the lines represents the principal component score for a given protein. **(B-C)** PCA feature plots of RPS13 and RPL38. **(D)** Unsupervised hierarchical clustering analysis of proteins constituting the cytosolic ribosome. **(E)** Structural model of the 80S ribosome (PDB: 4V6X) highlighting ribosomal proteins displaying variable abundance across skeletal muscle fibers. **(F)** Ribosomal proteins with variable stoichiometry located close to the mRNA exit tunnel.

The concept of ribosomal heterogeneity and specialization was previously introduced, where the presence of distinct subpopulations of ribosomes (ribosomal heterogeneity) can directly influence protein translation in different tissues^32^ and cells^33^ by selectively translating specific mRNA transcript pools^34^ (ribosomal specialization). To identify sub-sets of ribosomal proteins that are co-expressed within skeletal muscle fibers, we performed an unsupervised hierarchical clustering analysis of ribosomal proteins within the proteome (**Fig 3D, Table S10**). As expected, ribosomal proteins did not cluster by MYH-based fiber type. However, we identified three distinct clusters of ribosomal proteins; the first of which (ribosomal_cluster_1) was coregulated alongside RPL38, therefore elevated in fibers in the positive direction of PC1. The second cluster (ribosomal_cluster_2) was coregulated alongside RPS13 and was elevated in fibers in the negative direction of PC1. A third cluster (ribosomal_cluster_3) displayed no coordinated differential expression within skeletal muscle fibers and could therefore be considered “core” ribosomal proteins within skeletal muscle. Both ribosomal_cluster_1 and ribosomal_cluster_2 contain ribosomal proteins previously demonstrated to regulate selective translation (e.g. RPL10A, RPL38, RPS19 and RPS25) and functionally influence development (e.g. RPL10A, RPL38)^34–38^. In line with the results from the PCA analysis, the observed heterogeneous abundance of these ribosomal proteins across fibers was also of a continual nature (**Fig 3 S1C**).

To visualize the position of the ribosomal proteins that display heterogeneity within the ribosome, we utilized a structural model of the human 80S ribosome (Protein Data Bank: 4V6X)^39^ (**Fig 3E**). Upon highlighting the ribosomal proteins that belong to the different ribosomal clusters, they were not located in close proximity, showing that our approach did not enrich for a particular area/section of the ribosome. Yet interestingly, ribosomal_cluster_2 contained a lower proportion of large ribosomal subunit proteins than ribosomal_cluster_1 and ribosomal_cluster_3 (**Fig 3 S1D**). We observed that the majority of the proteins that display variable stoichiometry within skeletal muscle fibers are located on the surface of the ribosome (**Fig 3E**), which is consistent with an ability to interact with internal ribosome entry site (IRES) elements within distinct mRNA populations to coordinate selective translation^40,41^. Furthermore, numerous proteins that display variable stoichiometry within skeletal muscle fibers are located close to functional regions, such as the mRNA exit tunnel (**Fig 3F**), which can selectively regulate translation elongation and arrest of specific nascent peptides^42^. Overall, our data identifies that skeletal muscle ribosomal protein stoichiometry displays heterogeneity, driving variance between skeletal muscle fibers.

### Slow and fast fiber signatures and their transcriptional regulators

We next set out to identify novel features of fast and slow skeletal muscle fibers and how these are transcriptionally regulated. Comparing the fast and slow clusters defined in the UMAPs of both datasets (**Fig 1G-H & inlays Fig 4A-B**), transcriptome and proteome analysis yielded 1366 and 804 differentially abundant features, respectively (**Fig 4A-B, Table S11-14**). Expected differences in sarcomeric (e.g. tropomyosin and troponin), excitation-contraction coupling (SERCA isoforms) and energy metabolism-related (e.g. ALDOA and CKB) features were observed. In addition, transcripts and proteins regulating protein ubiquitination displayed differences between fast and slow fibers (e.g. *USP54, SH3RF2,* USP28 and USP48) (**Fig 4A-B**). Furthermore, the microprotein-encoding gene *RP11-451G4.2 (DWORF)*, which has previously been shown to be differentially expressed between lamb muscle fiber types^43^ and to enhance SERCA activity in cardiac muscle^44^, was significantly up-regulated in slow skeletal muscle fibers (**Fig 4A**). Also at the single fiber level, clear differences could be observed for known features such as the metabolism-related isoforms of lactate dehydrogenase (LDHA and LDHB, **Fig 4C** and **Fig 4 S1A**)^45,46^, as well as previously unknown fiber type-specific features (e.g. *IRX3*, *USP54*, USP28, and DPYSL3) (**Fig 4C**). There was a reasonable overlap of differentially expressed features (**Fig 4 S1B**) and correlation of fold changes between the transcriptomic and proteomic datasets, primarily driven by the large expression differences in sarcomeric features (**Fig 4 S1C**). Notably, some features (e.g. USP28, USP48, GOLGA4, AKAP13) showed strong post-transcriptional regulation with slow/fast fiber type-specific expression profiles only at the proteome level (**Fig 4 S1C**).

**Figure 4.**
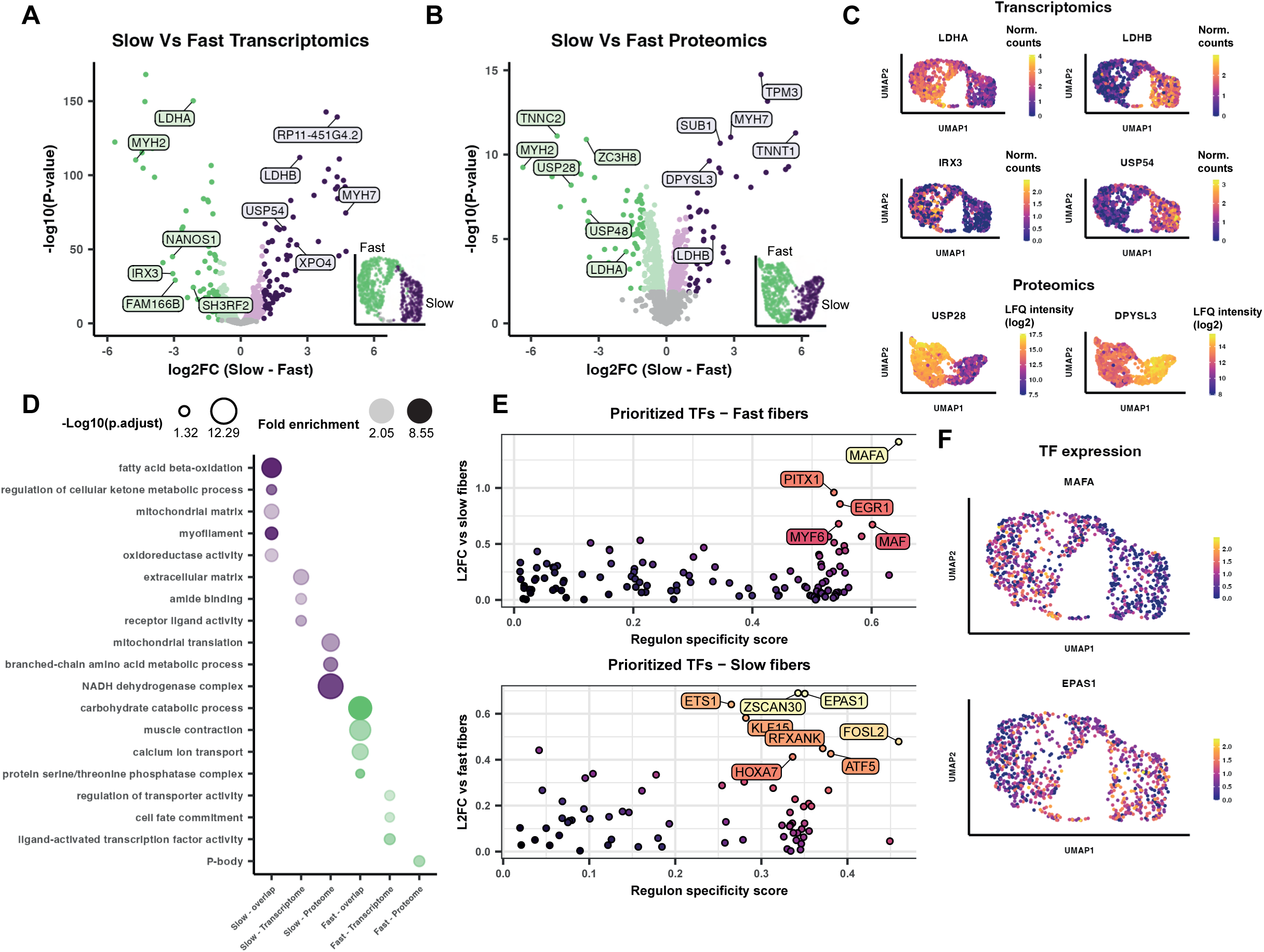
Fast and slow skeletal muscle fiber signatures. **(A-B)** Volcano plots comparing the slow and fast clusters identified from the uniform manifold approximation and projection (UMAP) plots in Fig 1G-H. Coloured dots represent significantly different transcripts or proteins at an FDR < 0.05, darker dots represent significantly different transcripts or proteins with a log fold change > 1. (**C**) Feature plots of selected differentially expressed genes or proteins between slow and fast fibers. **(D)** Enrichment analysis of significantly different transcript and proteins. Overlap – enriched in both datasets, Transcriptome – enriched only in transcriptome, Proteome – enriched only in proteome. **(E)** Fiber type specific transcription factors as identified by SCENIC, based on SCENIC-derived regulon specificity score and differential mRNA expression between fiber types. (**F**) Feature plots of selected differentially expressed transcription factors between slow and fast fibers.

We next performed over-representation analysis on the differentially abundant genes and proteins (**Fig 4D, Table S15**). Enriched pathways of features that were differential in both datasets showed expected differences such as fatty acid beta-oxidation and ketone metabolic process (slow fibers), myofilament/muscle contraction (fast and slow fibers, respectively) and carbohydrate catabolic process (fast fibers). Serine/Threonine protein phosphatase activity was also enriched in fast fibers, driven by features such as phosphatase regulatory and catalytic subunits (PPP3CB, PPP1R3D, and PPP1R3A), known to regulate glycogen metabolism^47^ (**Fig 4 S1D-E**). Other enriched pathways in fast fibers were Processing (P-) bodies (YTHDF3, TRIM21, LSM2) in the proteome (**Fig 4 S1F**), possibly related to post-transcriptional regulation^48^, and transcription factor activity (*SREBF1*, *RXRG*, *RORA*) in the transcriptome (**Fig 4 S1G**). Slow fibers showed enrichment for oxidoreductase activity (*BDH1*, *DCXR*, *TXN2*) (**Fig 4 S1H**), amide binding (*CPTP*, *PFDN2*, *CRYAB*) (**Fig 4 S1I**), extracellular matrix (*CTSD*, *ADAMTSL4*, *LAMC1*) (**Fig 4 S1J**) and receptor-ligand activity (*FNDC5*, *SPX*, *NENF*) (**Fig 4 S1K**).

To get more insights into the transcriptional regulation of slow/fast fiber type signatures, we performed transcription factor enrichment analysis using SCENIC^49^ (**Table S16**). Many transcription factors were significantly enriched between fast and slow fibers (**Fig 4E**). This included transcription factors such as *MAFA*, previously linked to the development of fast fibers^50^, but also multiple novel transcription factors potentially involved in the fiber type specific gene program. These included *PITX1*, *EGR1* and *MYF6* as the most enriched transcription factors within fast fibers (**Fig 4E**). Conversely, *ZSCAN30* and *EPAS1* (also known as *HIF2A*) were the most enriched transcription factors within slow fibers (**Fig 4E**). In line with this, *MAFA* expression levels were higher in the UMAP area corresponding to fast muscle fibers, whereas an opposite expression pattern was observed for *EPAS1* (**Fig 4F**).

### Non-coding RNA and associated microproteins in human skeletal muscle fibers

Alongside known protein-coding genes, there exists a multitude of non-coding RNA biotypes, potentially involved in the regulation of human development and disease^51,52^. Several non-coding RNAs displayed fiber type specificity in the transcriptomics dataset (**Fig 5A & Table S17**), including *LINC01405*, which is very specific to slow fibers and is reported to be downregulated in muscle of mitochondrial myopathy patients^53^. Conversely, *RP11-255P5.3,* corresponding to the *lnc-ERCC5-5* gene (https://lncipedia.org/db/transcript/lnc-ERCC5-5:2)^54^ displayed fast fiber type-specificity. Both *LINC01405* (https://tinyurl.com/x5k9wj3h) and *RP11-255P5.3* (https://tinyurl.com/29jmzder) display specificity to skeletal muscle (**Fig 5 S1A-B**) and have very few or no known contractile genes within a 1Mb genomic neighborhood, suggesting a specialized role in fiber type regulation rather than a regulatory role for a neighboring contractile gene. The slow/fast fiber type-specific specific expression profiles of *LINC01405* and *RP11-255P5.3* were independently validated using RNAscope (**Fig 5B-C)**.

**Figure 5.**
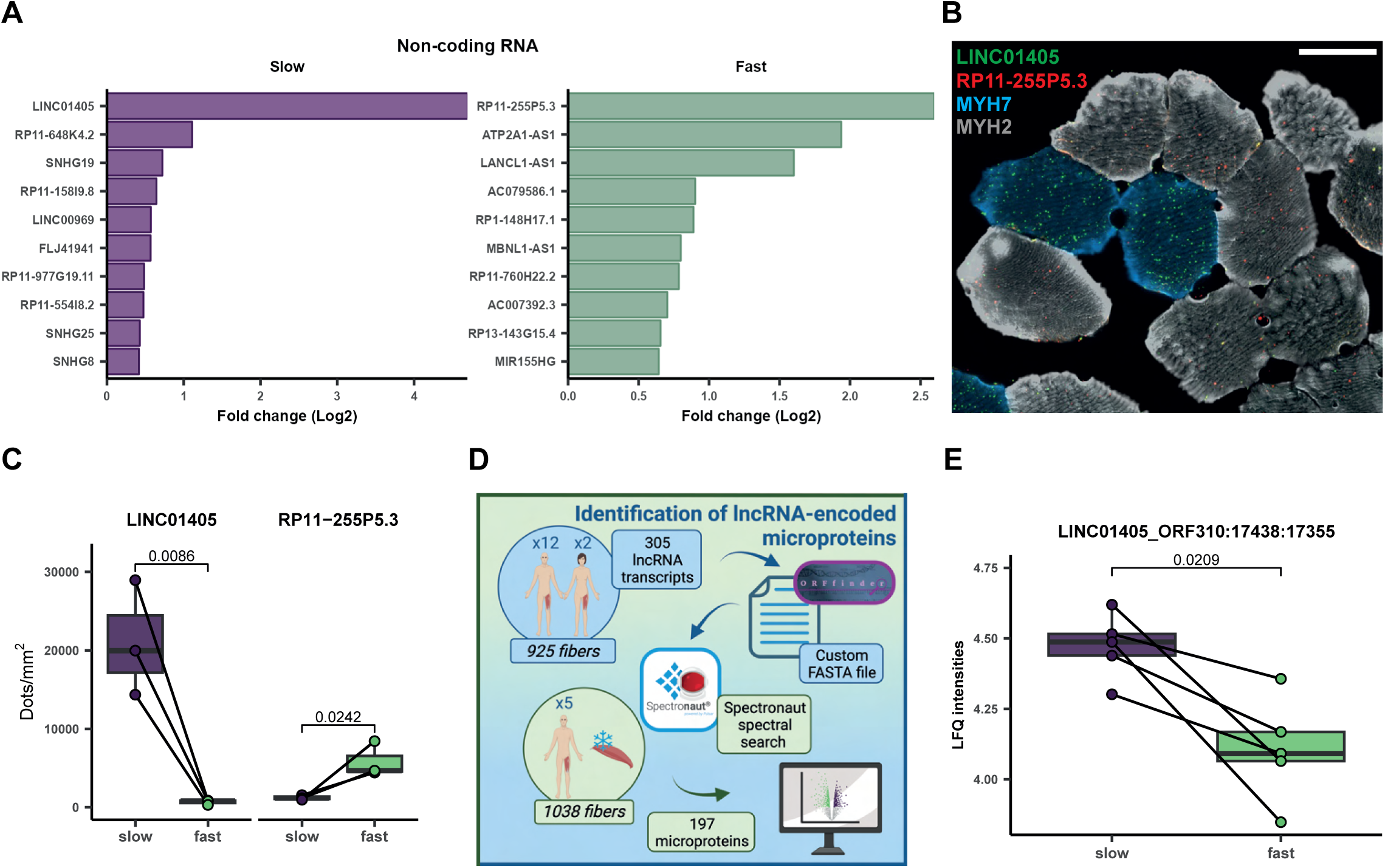
Non-coding RNA and associated microproteins in human skeletal muscle fibers. **(A)** Significantly regulated non-coding RNA transcripts in slow and fast fibers. **(B)** Representative RNAscope images displaying the slow/fast fiber-type specificity of LINC01405 and RP11-255P5.3, respectively. Scaling bar = 50 μm. **(C)** Quantification of RNAscope data for fiber type specific non-coding RNA expression. **(D)** De novo microprotein identification workflow. **(E)** The microprotein LINC01405_ORF408:17441:17358 is specific to slow skeletal muscle fibers.

Recently, it has become apparent that numerous assumed non-coding transcripts code for transcribed microproteins, some of which regulate muscle function^44,55^. To identify novel microproteins with potential fiber-type specificity, we searched our 1000 fiber proteomic dataset using a custom FASTA file containing sequences from the detected non-coding transcripts (n = 305) from the 1000 fiber transcriptome dataset (**Fig 5D**). This resulted in the identification of 197 novel microproteins arising from 22 distinct transcripts, of which 71 microproteins display differential regulation between slow and fast skeletal muscle fibers (**Fig 5 S1C and Table S18**). Three microprotein products were identified for *LINC01405*, of which one displays a slow-fiber type specificity similar to its transcript (**Fig 5E and 5 S1D**). Thus, we identify *LINC01405* as a microprotein-encoding gene, which displays specificity for slow skeletal muscle fibers.

### Nemaline myopathies induce a shift towards faster, less oxidative skeletal muscle fibers

Having developed a comprehensive workflow to characterize the proteome of single muscle fibers at scale and discovered novel regulators of fiber heterogeneity in the healthy state, we applied this pipeline to understand how nemaline myopathy impacts skeletal muscle fiber heterogeneity. Nemaline myopathy is a genetic muscular disorder that causes muscle weakness, resulting in a range of complications for the affected children including respiratory difficulties, scoliosis, and physical immobility^19,20^. Typically, in nemaline myopathy, pathogenic variants in genes such as actin alpha 1 (*ACTA1*) drive the fiber type composition towards slow fiber predominance, although this effect is heterogenous. The only clear exception is the troponin T1 (*TNNT1*) nemaline myopathy, in which a fast fiber predominance is seen. Thus, a deeper characterization of the heterogeneity behind the skeletal muscle fiber dysregulation observed in nemaline myopathies may help untangle the complex relationship between these diseases and muscle fiber types.

Muscle fibers isolated from patients with *ACTA1*- and *TNNT1*-mutation derived nemaline myopathies display substantial myofiber atrophy or hypotrophy, compared to healthy controls (n = 3 per group) (**Fig 6A, Table S19**), which provides considerable technical challenge arising from the limited material for proteomic analysis. Nonetheless, we were able to detect 2485 proteins from 272 skeletal muscle fibers. After filtering for a minimum of 1000 quantified proteins per fiber, downstream bioinformatic analyses were performed on 250 fibers. On average of 1573 ± 359 proteins were quantified per fiber after filtering (**Fig 6 S1A, Table S20-21**). Importantly, only a modest reduction in proteome depth was apparent in samples from patients with nemaline myopathy, despite the markedly reduced fiber size. Furthermore, processing of this data with our custom FASTA file (including non-coding transcripts) identified five microproteins within the skeletal muscle fibers from nemaline myopathy patients (**Table S22**). A wide dynamic range was apparent in the proteome, whilst the shared proteins in the control participants correlated well with those in the previous analysis of the 1000 fiber proteome study (**Fig 6 S1B-C**).

**Figure 6.**
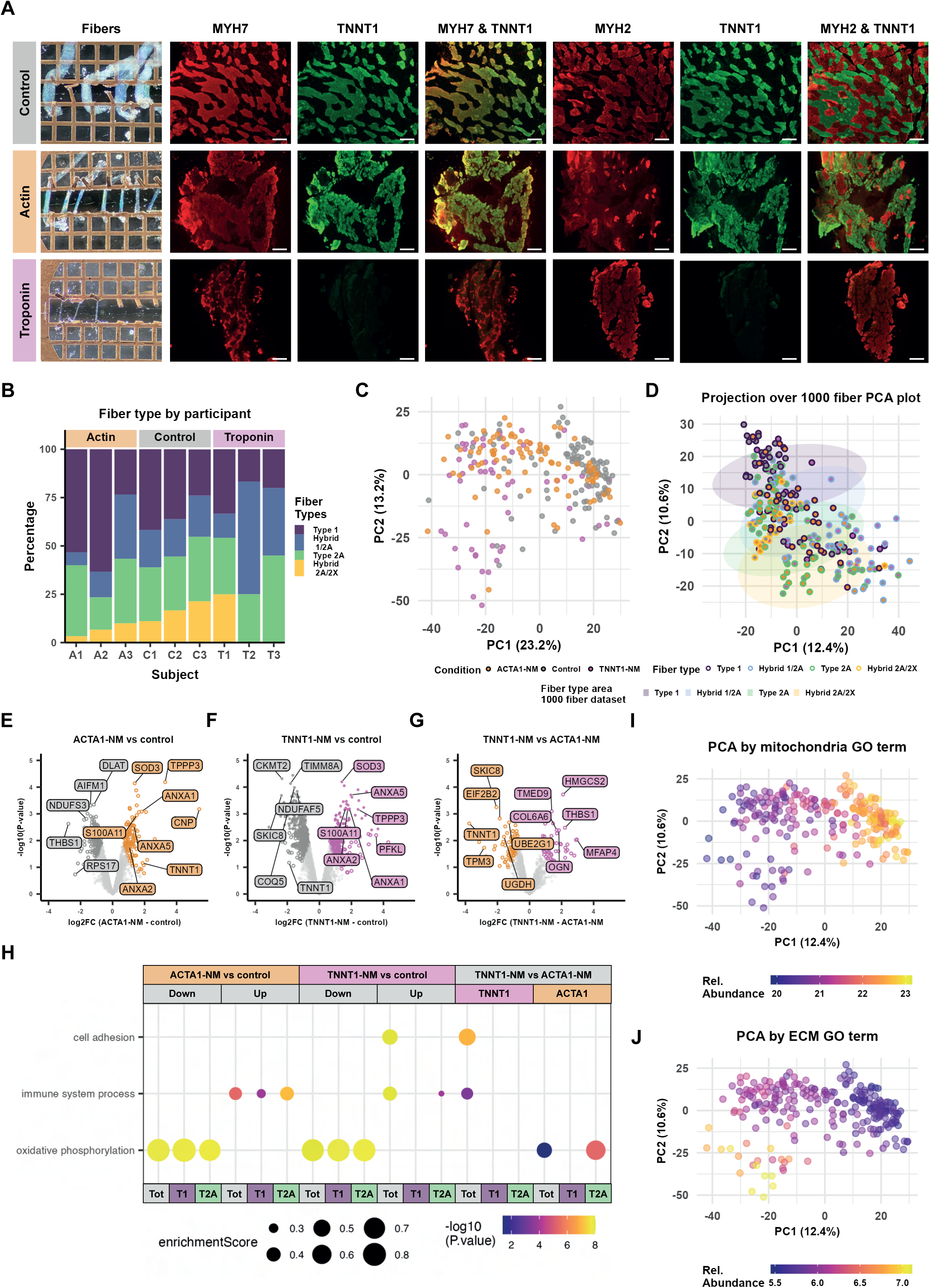
Nemaline myopathies induce a shift towards faster, less oxidative skeletal muscle fibers. **(A)** Microscopy images displaying fiber atrophy or hypotrophy and divergent MYH-based fiber type predominance in *ACTA1-* and *TNNT1-*nemaline myopathies (NM). Scaling bar = 100 μm. **(B)** MYH-based fiber type proportions by participant. **(C)** Principal component analysis (PCA) plot of skeletal muscle fibers from nemaline myopathy patients and controls. **(D)** Skeletal muscle fibers from nemaline myopathy patients and controls projected onto PCA plot determined from the 1000 fibers analyzed in Fig 2. **(E-G)** Volcano plots comparing the *ACTA1*- and *TNNT1*-nemaline myopathies with control participants and between *ACTA1*- and *TNNT1*-nemaline myopathies. Colored circles represent significantly different proteins at π < 0.05, darker dots represent significantly different proteins at FDR < 0.05. **(H)** Enrichment analysis of significantly different proteins in the whole proteome and within type 1 and type 2A fibers. **(I-J)** Principal component analysis (PCA) plot colored by extracellular matrix and mitochondria gene ontology (GO) terms.

As nemaline myopathies influence the MYH-based fiber type proportions within skeletal muscle^19,20^, we first investigated the MYH-based fiber type of our nemaline myopathy patients and controls. Fiber type was determined using the unbiased approach described previously for the 1000 fiber studies (**Fig 6 S1D-E**) and once again pure type 2X fibers could not be identified (**Fig 6B**). We observed a heterogeneous effect of nemaline myopathies on fiber type, as two patients with *ACTA1* mutations displaying an increased proportion of type 1 fibers, whilst two patients with *TNNT1*-nemaline myopathy displayed a reduced proportion of type 1 fibers (**Fig 6B**). Indeed, MYH2 and the fast troponin isoforms were downregulated in *ACTA1*-nemaline myopathy (TNNC2, TNNI2, and TNNT3), whilst MYH7 was downregulated in *TNNT1*-nemaline myopathy (**Fig 6 S2A**). This is in line with previous reports of heterogeneous fiber type switching in nemaline myopathies^19,20^. We validated these findings using immunohistochemistry, finding type 1 fiber predominance in *ACTA1*-nemaline myopathy patients, while the opposite was observed in *TNNT1-nemaline myopathy* patients (**Fig 6A**).

At the single-fiber proteome level, skeletal muscle fibers from *ACTA1*- and *TNNT1*-nemaline myopathy patients clustered away from the majority of control fibers, with TNNT1-nemaline myopathy fibers tending to be the most severely affected (**Fig 6C**). This was particularly apparent when we produced a PCA plot of pseudo-bulked fibers for each patient, with *TNNT1*-nemaline myopathy patients 2 & 3 lying furthest from the control samples (**Fig 6 S2B, Table S23**). To further understand how fibers from myopathy patients compare to the healthy condition, we capitalized on the depth of information provided by our analysis in 1000 fiber proteome study from healthy adult participants. We projected the fibers from our myopathy data set (both *ACTA1*- and *TNNT1*-nemaline myopathy patients and controls) onto the PCA determined from the 1000 fiber proteome study (**Fig 6D**). Control fibers displayed a similar distribution of MYH-based fiber type along PC2 as the 1000 fiber proteome study. However, the majority of fibers from nemaline myopathy patients shifted downwards along PC2, overlapping with the healthy fast fibers, irrespective of their own MYH-based fiber type. Thus, despite evidence for a fiber-type shift towards type 1 fibers in *ACTA1*-nemaline myopathy patients when quantified using MYH-based approaches, both *ACTA1*- and *TNNT1*-nemaline myopathies shift the skeletal muscle fiber proteome to resemble fast fibers.

We next directly compared each patient group with the samples from healthy controls, resulting in 256 and 552 differentially abundant proteins in *ACTA1*- and *TNNT1*-nemaline myopathies, respectively (**Fig 6E-G & Fig 6 S2C, Table S24**). Gene set enrichment analysis identified the coordinated reduction of mitochondrial proteins (**Fig 6H-I, Table S25**). Surprisingly, this was completely independent of MYH-based fiber type (**Fig 6H & Fig 6 S2D-I, Table S26**), despite the divergent fiber type predominance between *ACTA1*- and *TNNT1*-nemaline myopathies. Three microproteins were also regulated by either *ACTA1*- or *TNNT1*-nemaline myopathies. Two of those microproteins, ENSG00000215483_TR14_ORF67 (also known as *LINC00598* or *Lnc-FOXO1*) and ENSG00000229425_TR25_ORF40 (*lnc-NRIP1-2*) displayed differential abundance only within type 1 fibers, with

ENSG00000215483_TR14_ORF67 having previously reported to play a role in cell cycle regulation^56^. On the other hand, ENSG00000232046_TR1_ORF437 (corresponding to *LINC01798*) was upregulated in both type 1 and type 2 fibers from *ACTA1-*nemaline myopathy when compared to healthy controls (**Fig 6 S3A, Table S27**). Conversely, ribosomal proteins were largely unaffected by nemaline myopathies, although RPS17 was downregulated in *ACTA1-*nemaline myopathy (**Fig 6E**).

Enrichment analysis also identified the up-regulation of immune system process in both *ACTA1*- and *TNNT1*-nemaline myopathies, as well as cell adhesion in *TNNT1*-nemaline myopathy (**Fig 6H**). The enrichment of these extracellular terms, was reflected by extracellular matrix proteins driving the PCA in the negative direction in both PC1 and PC2, i.e. towards the most affected fibers (**Fig 6J**). Both patient groups overexpressed extracellular proteins involved in immune response and the sarcolemma repair machinery, such as annexins (ANXA1, ANXA2, ANXA5)^57,58^ and their interactor S100A11^59^ (**Fig 6 S3B-C**). This process has been previously reported to be upregulated in muscle dystrophies^60^, yet to our knowledge it has not been associated with nemaline myopathies before. A normal function of this molecular machinery is required for both sarcolemma membrane repair upon injury and for the fusion of new myoblasts to muscle fibers^58,61^. Therefore, an up-regulation of this process in both patient groups suggests a reparative response to damage caused by myofibril instability.

The effects of each nemaline myopathy were well correlated (r = 0.736) and displayed reasonable overlap (**Fig 6 S2A-B**), indicating that *ACTA1*- and *TNNT1*-nemaline myopathies induce similar effects on the proteome. Nonetheless, a number of proteins displayed regulation in only *ACTA1*- or *TNNT1*-nemaline myopathies (**Fig 6 S2A & S2C**). MFAP4, a pro-fibrotic protein, was one of the most up-regulated proteins in *TNNT1*-nemaline myopathy whilst unchanged in *ACTA1*-nemaline myopathy. SKIC8, a component of the PAF1C complex, which regulates the transcription of *HOX* genes^62^, was down-regulated in *TNNT1*-nemaline myopathy but was unaffected in *ACTA1*-nemaline myopathy (**Fig 6 S2A**). Direct comparisons between *ACTA1*- and *TNNT1*-nemaline myopathies identify a greater effect of *TNNT1*-nemaline myopathies on the reduction of mitochondrial proteins and the increase of immune system proteins (**Fig 6G-H & Fig 6 S2C & S2H-I**). These data are consistent with the greater degree of atrophy/hypotrophy apparent in *TNNT1*-compared to *ACTA1*-nemaline myopathies (**Fig 6A**), indicating that *TNNT1*-nemaline myopathy is the more severe form of the disease.

In order to assess whether the observed effects of nemaline myopathy were also present at the whole muscle-level, we conducted a bulk proteome analysis on muscle biopsies from the same *TNNT1-*nemaline myopathy patients and compared them against control individuals (n = 3 per group) **(Fig 6 S4A, Table S28)**. As expected, upon PCA, control individuals tightly clustered together whereas, similarly to the single fiber analysis, and *TNNT1-*nemaline myopathy patients displayed a higher inter-sample variance **(Fig 6 S4B)**. The bulk analysis was able to recapitulate the differentially expressed proteins **(Fig 6 S4C, Table S29)** and biological processes **(Fig 6 S4D, Table S30)** highlighted in the single fiber comparison, although lost the potential to discriminate between fiber types, and did not account for the heterogeneous effect of the disease across different fibers.

Together, these data demonstrate that single muscle fiber proteomics can illuminate clinical biology that is unapparent using targeted approaches such as immunoblotting. Furthermore, these data highlight the limitations of solely relying upon MYH-based fiber typing to describe phenotypic adaptations. Indeed, despite divergent MYH-based fiber type switching in actin and troponin nemaline myopathies, both nemaline myopathies uncouple the MYH-based fiber type from skeletal muscle fiber metabolism, shifting towards a faster, less oxidative muscle proteome.

## Discussion

Cellular heterogeneity is important to enable tissues to meet a wide range of demands. In skeletal muscle, this has classically been described by fiber types characterized by varying degrees of force production and fatigability. It is apparent, however, that this explains just a fraction of the variation in skeletal muscle fibers, with far greater variability, complexity and nuance residing in skeletal muscle fibers than previously believed. With the advent and development in technologies, these elements that regulate skeletal muscle fibers are now able to be elucidated. Indeed, our data indicates that type 2X fibers may not be a distinct sub-classification of skeletal muscle fibers. Furthermore, we identify metabolic, ribosomal, and cell junction proteins as major determinants of skeletal muscle fiber heterogeneity. By applying our proteomics workflow to samples from nemaline myopathy patients, we further evidence that MYH-based fiber typing does not capture the complete heterogeneity of skeletal muscle, particularly when the system is perturbed. Indeed, nemaline myopathies induce a shift towards faster less oxidative fibers regardless of their MYH-based fiber type.

Skeletal muscle fibers have been classified since the 19^th^ century^2,63^. Recent omics-based analyses have enabled us to start understanding expression profiles and responses to various stimuli specific to distinct MYH-based fiber types^11–16,24,46,64,65^. As demonstrated herein, omics approaches also have the advantage of increased sensitivity in the quantification of fiber-type markers over traditional antibody-based approaches, whilst also not relying on the quantification of a single (or few) markers for determining skeletal muscle fiber type. We leveraged complementary transcriptomics and proteomics workflows, and integrated the results, to explore the transcriptional and post-transcriptional regulation of human skeletal muscle fiber heterogeneity. This pipeline resulted in the observation that pure type 2X fibers could not be identified at the protein level in *vastus lateralis* skeletal muscle of our cohort of young healthy males. This is in line with previous single-fiber studies identifying <1% pure type 2x fibers in healthy *vastus lateralis*^66^, although this should be confirmed in other muscles in the future. The discrepancy between the identification of near pure type 2X fibers at the mRNA level but only hybrid type 2A/2X fibers at the protein level is puzzling. mRNA expression of MYH isoforms is not circadian^67^, indicating that it is unlikely that we have simply “missed” the on-signal of MYH2 in the apparently pure type 2X fibers at the RNA level. One possible explanation may be differences in the protein and/or mRNA stability of MYH isoforms, though this is purely hypothetical. Indeed, no fast fiber was 100% pure for any MYH isoform, though whether levels of MYH1 mRNA expression in the range of 70-90% could result in similar MYH1 and MYH2 abundance at the protein level is unclear. Nonetheless, when the whole transcriptome or proteome is considered, clustering-based analyses could only confidently identify two distinct clusters, which represented slow and fast skeletal muscle fibers regardless of their exact MYH composition. This is consistent with analyses using single nuclei transcriptomics approaches which commonly identify only two distinct clusters of myonuclei^68–70^. Furthermore, whilst previous proteomics-based studies have identified type 2X fibers, these fibers did not cluster away from the rest of the fast fibers and only displayed a handful of differentially abundant proteins compared to other MYH-based fiber types^14^.

These findings indicate that we should revert to the view of muscle fiber classification from the early twentieth century and classify human skeletal muscle fibers not into three distinct classifications based on MYHs, but instead into just two clusters based on their metabolic and contractile properties^63^.

Better still, we should consider muscle fiber heterogeneity in multiple dimensions. Previous omics studies already pointed in this direction, by showing that skeletal muscle fibers do not form discrete clusters, yet fall along a continuum^11,13,14,64,71^. Here, we found that over and above the variance within the contractile and metabolic signatures of skeletal muscle, fibers could also be separated by features related to the cell junction and translation machinery. Indeed, we identified ribosomal heterogeneity across skeletal muscle fibers that drives a slow/fast-fiber type independent heterogeneity. The underlying reason for such vast slow/fast-fiber type independent heterogeneity amongst fibers is not immediately obvious, though this could allude to a specialized spatial organization within a muscle fascicle to enable an optimal response to specific forces and loads^72^, specialized cellular or organ communication with other cell types within the muscle microenvironment^73–75^, or differential ribosomal activity in individual fibers. Indeed, ribosomal heterogeneity, via paralog substitution of RPL3 and RPL3L or at an rRNA 2ʹO-methyl level, has been implicated in skeletal muscle hypertrophy^76,77^. Multi-omics and spatial applications in concert with functional characteristics of single muscle fibers will further advance our understanding of muscle biology on a multi-omics level^78^.

In analyzing the proteome of single muscle fibers from patients with nemaline myopathies, we also demonstrate the utility, power, and applicability of single muscle fiber proteomics in uncovering clinical pathophysiology in skeletal muscle. Furthermore, by comparing our workflow against a bulk proteome analysis, we were able to demonstrate that single muscle fiber proteomics is able to capture the same depth of information than bulk tissue omics and expand on it by including inter-fiber heterogeneity and muscle fiber type into consideration. In addition to observing expected, albeit divergent, differences in fiber-type proportions in *ACTA1*- and *TNNT1*-nemaline myopathies compared to healthy controls^19^, we also identified oxidative and extracellular remodeling, which is uncoupled from this MYH-based fiber type switch. Fibrosis has previously been reported for *TNNT1* nemaline myopathy^19^. However, our analysis builds upon this to also identify the up-regulation of stress-related secreted proteins within the extracellular space, such as annexins, involved in the sarcolemma repair machinery^57–59^ in fibers from both *ACTA1*- and *TNNT1*-nemaline myopathy patients. Overall, the upregulation of annexins in muscle fibers from nemaline myopathy patients may represent a cellular response to rescue severely atrophying fibers.

Whilst this study represents the largest omic analysis of intact human single muscle fibers to date, it is not without limitations. We isolated skeletal muscle fibers from a relatively small and homogenous sample of participants and from a single muscle (*vastus lateralis*). In this respect, it is impossible to rule out the existence of specific fiber populations in different muscle types and at the extremes of muscle physiology. For example, we cannot rule out that a sub type of ultra-fast fibers (e.g. pure type 2X fibers) may become apparent in highly trained sprint and/or power athletes^79^ or during muscle disuse^66,80^. Furthermore, our limited participant pool did not allow us to investigate sex differences in fiber heterogeneity, in which there are known differences in fiber type proportions between males and females. Furthermore, we were unable to perform transcriptomics and proteomics on the same muscle fibers or in samples from the same participants. As we and others continue to optimize single-cell and single muscle fiber omics-analyses towards ultra-low sample input (as demonstrated here with the analysis of fibers from nemaline myopathy patients), the possibility of combining multi-omics (and functional) approaches in a single muscle fiber is tantalizingly close.

Collectively, our data identifies and explains transcriptional and post-transcriptional drivers of heterogeneity within skeletal muscle. In particular, we provide data to question long-standing dogmas within skeletal muscle physiology related to the classical definitions of MYH-based fiber-types. In doing so we hope to reignite the debate and ultimately redefine our understanding of skeletal muscle fiber classifications and heterogeneity.

## Methods

### 1000 fiber transcriptomics

#### Participant information

Fourteen participants (12 males / 2 females) volunteered to take part in this study, which was approved by the Ethical Committee of Ghent University Hospital (BC-10237), in agreement with the 2013 Declaration of Helsinki and registered on ClinicalTrials.gov (NCT05131555). General characteristics of the participants can be found in **Table S1**. After oral and written informed consent, participants were medically screened before final inclusion. Participants were young (22-42 years old), healthy (no diseases and non-smoking) and moderately physically active. Maximal oxygen uptake was determined as marker of physical fitness during a graded incremental cycling test, as previously described^81^.

#### Muscle biopsy collection

Muscle biopsies were collected in the rested and fasted state, on three different days, separated by 14 days. As these samples were collected as part of a larger study, on each of these days participants ingested a placebo (lactose), an H1-receptor antagonist (540 mg fexofenadine) or an H2-receptor antagonist (40 mg famotidine) 40 minutes before muscle biopsy collection. We have previously shown that these histamine receptor antagonists do not affect the resting skeletal muscle state^81^, and no clustering was apparent based on condition in our quality control plots (**Fig 1 S3E & Fig 2 S2B**). Dietary intake was standardized 48 hours before each experimental day (41.4 kcal/kg body weight, 5.1 g/kg body weight carbohydrates, 1.4 g/kg body weight protein and 1.6 g/kg body weight fat per day), followed by a standardized breakfast on the morning of the experimental day (1.5 g/kg bodyweight carbohydrates). Muscle biopsies of the *m. vastus lateralis* were then collected after local anaesthesia (0.5 mL of 1% Xylocaïne without epinephrine) using the percutaneous Bergström technique with suction^82^. The muscle samples were immediately submerged in RNA*later* and stored at 4 °C until manual fiber dissection (max. 3 days).

#### Single fiber isolation

Freshly excised muscle fiber bundles were transferred to fresh RNA*later* in a petri dish. Individual muscle fibers were then manually dissected using a stereomicroscope and fine forceps. Twenty-five fibers were dissected per biopsy, with special care to select fibers from different sections of the biopsy. After dissection, each fiber was carefully submerged in 3 µL of lysis buffer (SingleShot Cell Lysis kit, Bio-rad), containing proteinase K and DNase enzymes to remove unwanted proteins and DNA. Next, cell lysis and protein/DNA removal was initiated by short vortexing, spinning liquid down in a microcentrifuge and incubation at room temperature (10 min). Lysates were then incubated for 5 min at 37 °C and 5 min at 75 °C in a thermocycler (T100, Bio-Rad), immediately followed by storage at -80 °C until further processing.

#### Sequencing library preparation

Illumina-compatible libraries from polyadenylated RNA were prepared from 2 µL of the muscle fiber lysates using the QuantSeq-Pool 3’ mRNA-Seq library prep kit (Lexogen). Detailed methodology can be found in the manufacturer guidelines. The process was initiated by reverse transcription for first strand cDNA synthesis, during which Unique Molecular Identifiers (UMIs) and sample-specific i1 barcodes were introduced enabling sample pooling and reducing technical variability in the downstream process. Next, cDNA from 96 fibers was pooled and purified using magnetic beads, followed by RNA removal and second strand synthesis by random priming. Libraries were purified using magnetic beads, followed by addition of pool-specific i5/i7 indices and PCR amplification. A last purification step was performed, finalizing the Illumina-compatible libraries. A high sensitivity small DNA Fragment Analysis kit (Agilent Technologies, DNF-477-0500) was used to assess the quality of each library pool.

#### Illumina sequencing

The individual pools were further equimolarly (2 nM) pooled, based on Qubit-quantified concentrations. The final pool was subsequently sequenced with a NovaSeq S2 kit (1 x 100 nucleotides) with a loading of 2 nM (4% PhiX) in standard mode on a NovaSeq 6000 instrument.

#### Primary data processing

Our pipeline was based on the QuantSeq Pool data analysis pipeline from Lexogen (https://github.com/Lexogen-Tools/quantseqpool_analysis). First, the data was demultiplexed based on the i7/i5 indices with bcl2fastq2 (v2.20.0). The next demultiplexing step was performed via idemux (v0.1.6) according to the i1 sample-specific barcodes in read 2, followed by extraction of UMI sequences with umi_tools (v1.0.1). Trimming of the reads was then performed in multiple rounds with cutadapt (v3.4), with removal of too short reads (length < 20) or reads consisting entirely of adapter sequences. Reads were then aligned to the human genome with STAR (v2.6.0c), followed by BAM file indexing with SAMtools (v1.11). Duplicates of reads were removed with umi_tools (v1.0.1). Finally, counting of the alignments was performed with featureCounts from Subread (v2.0.3). At several intermediate steps during the pipeline, quality control with performed with FastQC (v0.11.9).

#### Initial Seurat processing

All further bioinformatics processing and visualization was performed in R (v4.2.3), primarily with the Seurat v4 workflow^83^. The individual UMI counts and metadata matrices were thus transformed into a Seurat object. Genes with expression in less than 30% of all fibers were removed. Low-quality samples were then removed based on a minimum threshold of 1000 UMI counts and 1000 detected genes. This resulted in a total of 925 fibers that passed all quality control filtering steps. Normalization of UMI counts was performed using the SCTransform v2 Seurat method^84^, including all 7418 detected features and regressing out participant variation. All relevant metadata can be found in **Table S31**.

## Proteomics

### Sample collection - 1000 fiber proteome

#### Participant information

Stored biobank muscle specimens were used for the purpose of the present study (Clinicaltrials.gov identifier: NCT04048993). The specimens were collected from five active and healthy male volunteers (aged 21-35 years) who gave their written and oral informed consent with approval from the Science Ethics Committee of the Capital Region in Denmark (H-1-2012-090) and complied with the guidelines of the 2013 Declaration of Helsinki. General characteristics of the participants can be found in **Table S1.** Participants were young, healthy (no diseases and non-smoking) and moderately physically active.

#### Muscle biopsy collection

Participants arrived in the morning after an overnight fast and rested in the supine position for 1 hour. Then, local anesthesia (2-3 mL Xylocaine 2%; lidocaine without epinephrine, AstraZeneca, Denmark) was applied under the skin above the fascia at the belly of the *m. vastus lateralis* muscle. A muscle biopsy was sampled through a small 3-4 mm incision using a Bergström needle with suction. The muscle biopsy specimen was snap-frozen in liquid nitrogen and stored at −80°C until analysis.

### Single muscle fiber isolation

Muscle fibers were isolated from freeze-dried specimens as previously described^24^. In brief, muscle biopsies were freeze-dried for 48 hours. Subsequently, fibers were isolated in a humidity- and temperature-controlled room (humidity of 25%) using fine forceps under a stereomicroscope. Approximately 200 single muscle fibers were isolated for each biopsy, resulting in a total of 1038. To ensure the fibers settled at the bottom of the tube, each fiber-containing tube underwent centrifugation at 20,000 g using a small centrifuge. Next, fibers were resuspended in 15 µL of lysis buffer (1% sodium dodecyl sulfate (SDC), 40 mM chloroacetamide (CAA), 10 mM dithiothreitol (DTT) in 50 mM Tris pH 8.5).

### Sample collection – myopathy

#### Participant information

Six patients with severe nemaline myopathy were selected from our nemaline myopathy study cohort. Three patients had pathogenic variants in *ACTA1*, representing the conventional severe form, and three patients had pathogenic variants in *TNNT1*, resulting in a rare, progressive form of nemaline myopathy. Three healthy individuals with no history of neuromuscular disease were used as controls (**Table S19**).

#### Muscle biopsy collection

Healthy control participant biopsies (n = 3) were used from an original study^85^, and were therefore collected, snap frozen in liquid nitrogen and stored in - 80℃ as per that originally described. For the present study, a fragment of this stored biopsy was dissected under sterile, frozen conditions before being prepared for single myofiber isolation (detailed below). Acquisition of biopsies of healthy control patients was approved by local ethics committee (Copenhagen and Frederiksberg) in Denmark (hs:h-15002266). Those of myopathy patients were consented, stored, and used in accordance with the Human Tissue Act under local ethical approval in United Kingdom (REC 13/NE/0373). All procedures were carried out in accordance to the Declaration of Helsinki.

#### Single fiber isolation

Dissected fragments of muscle biopsy were placed in ice-cold, 22 micron filtered relaxing solution (4 mM Mg-ATP, 1 mM free Mg ^2+^, 10^-.00^ mM free Ca^2+^, 20 mM imidazole, 7 mM EGTA, 14.5 mM creatine phosphate, KCl to an iconic strength of 180 mM and pH to 7.0)^86^ for ∼3 minutes before being immersed in fresh relaxing solution on a sterile petri dish and mounted on ice under a dissection microscope for single fiber isolation. Fibers were cleaved from the tissue/biopsy ensuring a variety of sample/biopsy locations were used and that only single fibers were selected. Following isolation, fibers were manually moved to a sterile 96-well plate containing 15 µL of lysis buffer (identical to that detailed above) where the tweezers containing the fiber were submerged, spun and agitated in lysis buffer to ensure the fibers dissociated from the tweezers. To ensure fibers were settled to the bottom of the well, the 96-well plate was subjected to gentle vortexing and centrifugation (1,000 g).

### Proteomics analysis

#### Sample preparation

Samples from both proteomics studies followed the same sample preparation workflow. In order to extract the proteins, samples were boiled at 95°C in a thermomixer with gentle shaking (800 rpm) and sonicated in a bioruptor instrument with 30 seconds on/off cycles for 15 minutes. A small 5 µL fraction of lysate from each sample was saved for antibody-based fiber typing of the 1000 fiber samples. Next, samples were processed following a modified version of the in-solution digestion sample preparation protocol. In brief, total volume was adjusted to 50 µl by addition of digestion buffer, containing 50 mM Tris PH 8.5 buffer, an enzyme to protein ratio of 1:500 LysC (Wako) and a 1:100 enzyme to protein ratio of trypsin (Promega). SMF lysates were digested overnight in a thermomixer set to 37⁰ C and 800 rpm. The next day, protein digestion was quenched by addition of 50 µl of 2% trifluoroacetic acid (TFA) in isopropanol. Peptides were desalted using in-house prepared single-use reverse-phase StageTips containing styrenedivinylbenzene reverse-phase sulfonate (SDB-RPS) disks. Then, desalted peptides were loaded in Evotips (Evosep) following manufacturer instructions prior LC-MS/MS analysis. Bulk tissue samples were prepared using the same protocol utilized for single fibers, with a few modifications to sample lysis. Tissue samples were first powdered using a tissue crusher over dry ice before resuspending the powder in the same lysis buffer described above. Then, the samples were homogenized using an IKA Turrax homogenizer for 2 minutes prior boiling and sonication. From here onwards the samples underwent the same protocol described above.

#### Proteomics library preparation

Fibers from the five healthy control individuals participating in the 1000 fiber study were carefully dissected and combined in order to create a pooled fiber lysate. Then, 200 µg of protein corresponding to each participant-specific lysate were pooled together into one final protein lysate that was processed following the same sample preparation workflow just described. 20 µg of desalted peptides were fractionated using High pH Reverse-Phase Chromatography (HpH-RP). Fractionation was carried out on a Kinetex 2.6 µm EVO C18 100 Å, 150 x 0.3 mm column manufactured by Phenomenex and using an EASY-nLC 1200 System (Thermo) operating at 1.5 µL/min. Separation was accomplished using a 62 min step-gradient starting from 3% to 60% solvent B (which consisted of 10 mM TEAB in 80% acetonitrile) and solvent A (containing 10 mM TEAB in water). The total run time was 98 min, which included wash and column equilibration. Throughout the fractionation, peptides were eluted and collected every 60 s, obtaining 96 single fractions without concatenation. Finally. 200 ng of HpH-RP fractionated peptides were loaded, concentrated and desalted on Evotips (Evosep) following the instructions provided by the manufacturer.

#### Liquid chromatography tandem mass spectrometry

Proteomics measurements were performed using LC-MS instrumentation consisting of an Evosep One HPLC system (Evosep)^22^ coupled via electrospray ionization to a timsTOF SCP mass spectrometer (Bruker). Peptides were separated utilizing 8 cm, 150 μM ID columns packed with C18 beads (1.5 μm) (Evosep). Chromatographic separation was achieved by the ‘60 samples per day’ method, followed by electrospray ionization through a CaptiveSpray ion source and a 10 μm emitter into the MS instrument. Single muscle fiber peptides were measured in DIA-PASEF mode following a previously described method^23^, while library fractions were measured using DDA-PASEF. In brief, the DDA-PASEF scan range encompassed 100 - 1700 m/z for both MS and MS/MS, and TIMS mobility range was set to 0.6 – 1.6 (V cm^−2^). Both TIMS ramp and accumulation times were configured to 100 ms, and 10 PASEF ramps were recorded for a total cycle time of 1.17 s. The MS/MS target intensity and intensity threshold were defined as 20.000 and 1.000, respectively. An exclusion list of 0.4 min for precursors within 0.015 m/z and 0.015 V cm^−2^ width was also activated. For DIA-PASEF the scan range was established at 400-1000 (m/z), the TIMS mobility range to 0.64-1.37 (V cm^−2^), and ramp and accumulation times were both set to 100 ms. A short-gradient method was used, which included 8 DIA-PASEF scans with three 25 Da windows per ramp, resulting in an estimated cycle time of 0.95 sec.

#### MS data processing

Library files were processed using the MS-Fragger functionality within Fragpipe v19.0 under the SpecLib workflow with default settings, including a minimum peptide length of seven amino acids and maximum of two missed cleavages allowed^87^. Spectra were searched against a human reviewed FASTA from Uniprot (March 2022, 20410 entries) and the output library contained a total of 5350 protein groups and 84383 precursors. Sample raw MS files were analyzed using DIA-NN version 1.8^88^ in a library-based manner against the MS library just described. Protein groups quantification was based on proteotypic peptides, neural network was set to double-pass mode, quantification strategy was set to “Robust LC (high accuracy)” and the match between runs options was enabled, the rest of parameters remained as default, which included precursor FDR set to 1% and peptide length of 7-30 amino acids.

#### Data processing

Further data analysis was performed under the R environment (R version 4.22). Both a metadata data frame containing sample and participant information and the “PG_matrix.tsv” file from DIA-NN’s output were loaded in RStudio. The 1000 fiber data frame was filtered to remove samples with less than 50% valid protein intensity values, resulting in a total of 974 fibers. Next, rows were filtered to remove proteins with less than 30% valid values across samples, resulting in a total of 1685 proteins. Regarding the myopathy dataset, after filtering samples for 50% valid values, the number of samples was 250. We included in the analysis proteins that were quantified in 70% of the samples in at least one condition (conditions: control, actin myopathy and troponin myopathy), resulting in a total of 1545 proteins. Both data frames were then log2 transformed and normalized using the *normalizeBetweenArrays()* function from the limma package, with the method argument set to quantile^89^. Then, batch correction through the *ComBat()* function from the sva package^90^ was applied to minimize the effect of the three technical batches originated during mass spectrometry measurement. Finally, missing values were replaced by random numbers from a Gaussian distribution with the default settings of the *tImpute()* function from the PhosR package^91^. All relevant metadata for the 1000 fiber proteome and nemaline myopathy datasets can be found in **Tables S32 and S33, respectively**.

### Bioinformatics analysis

#### Transcriptome and proteome dynamic range

The expression/intensity for each gene/protein was calculated relative to the total counts/intensity for each fiber. This value was then averaged across fibers in each dataset and log10-transformed. The overlap of detected features between both datasets were analyzed using the *VennDiagram* package (v 1.7.3).

#### Coefficient of variation – proteomics

For each of the 96 well plates used during the MS measurement of the 1000 fiber study, one technical control sample was included in A1 position to monitor total ion current intensity and quality control of the runs (a total of eleven technical controls). The coefficient of variation between proteins was calculated by dividing the standard deviation by the mean of the LFQ intensities from each protein across technical replicates and then multiplied by one hundred.

#### Correlation analyses

Mean log2 transformed transcript counts, protein intensities, and/or fold change values across fibers were calculated, filtered for shared proteins/genes and Pearson correlation was calculated.

#### Omics-based fiber typing

Normalized counts and raw LFQ intensities were retrieved from well-described contractile proteins that have a slow (MYH7, TNNT1, TPM3, ATP2A2 and MYL3) and fast (MYH2, MYH1, TNNT3, TPM1, ATP2A1 and MYL1) isoform^2^. For each isoform combination, the relative expression of each was then calculated and samples were ordered from high to low. The mathematical bottom knee for each curve was then determined using the *barcodeRanks()* function in the DropletUtils package (v 1.18.1). This threshold was used to assign fiber types as pure (type 1, type 2A or type 2X) or hybrid (hybrid 1/2A, hybrid 2A/2X or hybrid 1/2X) (**Table S6**). For the features with only two isoforms, fibers were assigned as ‘slow’, ‘fast’ or ‘hybrid’. To determine the overlap of the contractile features assigning a fiber as being slow, upset plots were generated using the *upset()* function of the ComplexUpset package (v 1.3.3), and then simplified to bar plots.

#### Principal Component Analysis (PCA)

PCA was performed using the *RunPCA()* function of the Seurat package. Scree plots were based on the *fviz_eig()* function of the factoextra package (v 1.0.7), which worked after PCA with *prcomp()*.

#### Seurat clustering

Uniform Manifold Approximation and Projection (UMAP) clustering was performed based on the K-nearest neighbor graph with the first 6 dimensions as input for both the transcriptome and proteome datasets (**Fig 1 S3D-E**). Feature plots were generated using the *FeaturePlot()* function. UMAP plots were colored based on different criteria (MYH-based fiber types, participant, test day) stored in the metadata.

#### Enrichment analysis

Genes and protein sets were processed to obtain lists of features that were differentially expressed or, in the case of the top PCA drivers, in the top 5% drivers for the positive and negative direction of the first and second principal component. Over-representation analysis was then performed on these features with the *enrichGO()* and *simplify()* function of the clusterProfiler package (v 4.6.2)^92^ using all gene ontology terms. Obtained lists of significant terms were manually curated to extract interesting and relevant terms.

#### Hierarchical clustering of ribosomal proteins

Raw proteomics data was log2 transformed and filtered to contain proteins enlisted in the ‘cytosolic ribosome’ GO term, followed by Z-scoring prior heatmap visualization using the *pheatmap()* function from the Pheatmap package (v 1.0.12). The number of clusters was determined by visual inspection and assigning a value of 3 to the *cuttree*() function.

#### Differential expression analysis

To avoid artificially inflated p-values, which would arise from regarding every fiber as an independent replicate, we employed a pseudobulk differential expression analysis approach. We mathematically downsampled the total data points to one value per MYH-based fiber type per participant by aggregating (transcriptomics) or taking the median value (proteomics). Transcriptomics data were further processed using the DESeq2 pipeline (v 1.38.3)^93^ with a ‘∼ participant + fiber type’ statistical model. 1000 fiber proteomics data was processed using the limma (v 3.54.2) workflow, fitting the data to a linear model defined as: ‘∼ 0 + fiber type + participant‘, whereas the myopathy dataset was fitted to ‘∼ 0 + condition’ for the comparisons between conditions and ‘∼ 0 + fiber type and condition’ for the comparisons including fiber type. Fitted models were then subjected to gene ranking using an empirical Bayes method using *eBayes*() prior extracting the results through *topTable*(), with p-value adjustment set to Benjamini-Hochberg, both functions from the limma package. Threshold for significantly different genes/proteins was defined as adjusted p-values smaller than 0.05 and a log fold change cut-off of 1 was applied. For the nemaline myopathy dataset, the Xiao significance score was applied, which combines expression fold change and statistical significance^94^. Proteins with a Xiao score under 0.05 were regarded as differentially expressed between conditions.

#### SCENIC

Inference of active transcription factors in slow and fast fibers was performed using Single-Cell rEgulatory Network Inference and Clustering (SCENIC, pySCENIC version 0.12.1 with cisTarget v10 databases and annotations)^95^. To prioritize fiber-type specific transcription factors, both their fiber-type specific expression at mRNA level and regulon activity were combined into a final prioritization score. This prioritization score was calculated as the sum of the z-score scaled differential expression score (logFC from pseudobulked data) and z-score scaled regulon specificity scores (RSS).

#### Non-coding RNA

For the transcriptomics data, the biotype of each gene was determined using the ‘GENEBIOTYPE’ column using the AnnotationDbi package (v 1.60.2) with the EnsDb.Hsapiens.v86 database. Genomic location interrogation was performed using the UCSC Human Genome Browser (https://genome.ucsc.edu). Tissue-specific gene expression of interesting long non-coding RNAs was explored using the GTEx Portal database.

### Microprotein identification

#### Construction of putative lncRNA-encoded protein database

RNA sequences of the non-coding transcripts were extracted using the *getSequence() function from the biomaRt package* (v 2.56.1), with ‘transcript_exon_intron’ as the *seqType.* Both intergenic (lincRNA) and antisense long non-coding RNA (lncRNA) transcripts were utilized for database construction. A six-frame-translation was used to translate the corresponding RNA sequences into the proteins, as well as ORFfinder NCBI functionality (https://www.ncbi.nlm.nih.gov/orffinder/ ) was used to extract open reading frames (ORFs) from the transcripts. Minimal ORF length was set to 75 nucleotides, genetic code was set to “Standard” and “Any sense codon” was used, as a start codon, to extract maximum number of open reading frames. The obtained protein fasta file contained multiple entries for each gene name, ensured by various combinations of: i) transcript identifiers, ii) ORF identifiers and iii) start:stop codons.

#### Identification of lncRNA-encoded proteins

DIA raw MS data were analyzed with Spectronaut v18 using an in-house generated sample-specific fasta, comprised of the reviewed human proteome (proteome ID: UP000005640, 20 426 proteins, downloaded Sep 2023) and lncRNA-encoded protein sequences (125 223 proteins), in directDIA mode. The default settings were used unless otherwise noted. Data filtering was set to “Qvalue”. False discovery rate (FDR) was set to 1% at peptide precursor level and 1% at protein level. Top3 peptide precursors were used for protein quantification. The downstream data analysis was performed using in-house developed R script.

##### PCA projection

The 1000 fiber and myopathy data sets were initially filtered to remove non-overlapping proteins. Then, they were combined and normalized using the *normalizeBetweenArrays()* function from the limma package. The normalization method used was “quantile” to ensure that both data sets had the same distributions and were comparable. Subsequently, the merged data set was divided back into the two separate data sets, namely the 1000 fiber data set and the myopathy data set. For the 1000 fiber data set, PCA was calculated using the *prcomp()* function. Moving on, the myopathy data set was multiplied by the PC loadings obtained from the 1000 fiber dataset to generate its PCA projection. Finally, the PCA projections from the myopathy samples were plotted on the top of the 1000 fiber PCA visualization.

### Muscle-specific ribosomal gene signature

The skeletal muscle-specific ribosomal gene signature, consisting of log2 fold change values comparing the mRNA expression of ribosomal subunits in skeletal muscle compared to 52 other human tissues, was downloaded from Panda et al^96^. Log2 fold change values were ranked to identify ribosomal proteins with the highest overexpression in human skeletal muscle.

### Structural Analysis of Ribosomal Proteins

The human 80S ribosome structural model (Protein Data Bank: 4V6X) was downloaded from the Protein Data Bank website (RCSB PDB). Visualization and editing of the ribosomal structure, and preparation of figures and movies were performed in UCSF ChimeraX^97^.

### Antibody-based fiber typing

Dot blot was conducted following a previously described protocol with a few modifications^24^. Initially, two identical PVDF membranes were activated using 96% ethanol and washed with transfer buffer. Subsequently, the membranes were placed on wet filter paper with transfer buffer until they dried. Next, 1 µL of fiber lysate was spotted at the same position on both membranes, and the membranes were allowed to dry. Reactivation of the membranes was carried out using 96% ethanol, followed by gentle washing with TBST. The membranes were then blocked in TBST containing 5% skim milk for 15 minutes. After blocking, the membranes were washed three times with TBST and incubated with the primary antibody solution of either anti-MYH7 (A4.840) or anti-MYH2 (A4.74), both from Developmental Studies Hybridoma Bank (DSHB) at a dilution of 1:200 in TBST containing 1% skim milk for one hour. Subsequently, the membranes were gently washed three times with TBST and incubated with the secondary antibody (anti-mouse) at a dilution of 1:20,000 in TBST containing 1% skim milk for two hours. Finally, the membranes were washed three times for five minutes each with TBST and visualized using Immobilon Forte (Milipore) in a ChemiDoc XRS+ (Bio-Rad) imaging system.

### RNAscope

8µm sections from three different fixed-frozen human muscle biopsies were used for RNAscope labelling and subsequent immunohistochemistry (IHC). For detection of RP11-255P5.3 and LINC01405 commercially available RNAscope Multiplex Fluorescent Assay V2 (Advanced Cell Diagnostics) and probes against Lnc-ERCC5-5-C1 (# 1276851-C1), and LINC01405-C2 (# 549201-C2) (Advanced Cell Diagnostics), were used according to the manufacturer’s protocols. To control tissue quality Positive 3-plex probe (# 320881) and negative 3-plex probe (# 320871) were used. To visualize different subtypes of muscle fibers, sections after RNAscope were blocked with 5% donkey serum and incubated with antibodies against MYH2 (A4.74-s (1:5); DSHB) and MYH7 (A4.840 (1:5); BSHD) overnight. After washing sections were incubated with Alexa Flour 488-conjugated Donkey Anti-Mouse IgG, Fcγ Subclass 1 and DyLightTM405-conjugated Donkey Anti-Mouse IgM secondary antibodies respectively and mounted with ProLong™ Diamond Antifade Mountant (Invitrogen). Slides were imaged using Zeiss Axio Observer microscope equipped with Axiocam 702 camera. Biopsies from three individuals were used for quantification, with 187 muscle fibers being counted in total. As in RNAscope each dot corresponds to one RNA molecule, the number of dots/mm^2^ was used as a measure of RNA expression. We first determined the number of dots/mm^2^ within each fiber for both probes, then averaged the results by fiber type and participant. The given average was then used as an input for a two sample t-test.

### Immunostaining for muscle sections

Immunolabelling was performed on 10 μm cryosections, fixed in 4% PFA (10 min), permeabilized in 0.1% Triton X-100 (20 min) and blocked in 10% Normal Goat Serum (50062Z, Life Technologies) with 0.1% BSA (1 h). Sections were incubated o/n (4°C) with primary antibodies against MYH7 (mouse monoclonal A4.951, Santa Cruz, sc-53090, diluted 1:25) or MYH2 (mouse monoclonal SC71, DSHB, 1:25), each combined with primary antibody against TNNT1 (rabbit polyclonal HPA058448, Sigma, diluted 1:500) in 5% goat serum with 0.1% BSA and 0.1% Triton X-100. Alexa Fluor Goat anti-Mouse 647 (A21237) was used as the secondary antibody for the MYHs and Alexa Fluor Donkey anti-Rabbit 488 (A11034) for the TNNT1 (Life Technologies, 1:500 each in 10% Normal Goat Serum). Fluorescent images were obtained with a 10x objective on a Zeiss Axio Observer 3 fluorescence microscope with a Colibri 5 led detector, combined with Zeiss Axiocam 705 mono camera, using Zen software (Zeiss). For visualization purposes, a selection of fibres were mounted on copper grids glued on a microscopy slide, and imaged under a stereomicroscope.

## Data availability

RNA-seq data has been deposited at the European Genome-phenome Archive (EGA), which is hosted by the EBI and the CRG, under study ID EGAS50000000418. Raw mass spectrometry files and search engine results have been deposited at the ProteomeXchange Consortium through the PRIDE partner repository (identifier PXD046863).

## Code availability

All code used for this manuscript is available upon request.

## Acknowledgments

This work is supported by an unconditional donation from the Novo Nordisk Foundation (NNF) to NNF Center for Basic Metabolic Research (Grant numbers NNF18CC0034900 and NNF23SA0084103). Mass spectrometry analyses were performed by the Proteomics Research Infrastructure (PRI) at the University of Copenhagen (UCPH), supported by the Novo Nordisk Foundation (NNF) (grant agreement number NNF19SA0059305). This work was generously funded by grants from the Novo Nordisk Foundation (NNF-0070539; to JO), the Foundation Building Strength for Nemaline Myopathy (to JO) and the Research Foundation Flanders (G080321N, 11B4220N and 1268023N; to WD, TVdS and EL, respectively). Several plots were made with BioRender.com. The authors would like to thank Alberto Santos, Leonidas Lundell, and Lars Roed Ingerslev for their input on data analysis and visualization.

## Author contributions

Conceptualization: RMJ, TVdS, BS, EL, PM, JV, JO, MH, WD, ASD

Methodology: RMJ, TVdS, BS, JL, RAS, DS, OD, AVdL, EL, LMA, KVV, LM, MW, PM, JV, MH, WD, ASD

Software: RMJ, TVdS, BS, DS, LMA, RB, JA

Validation: RMJ, TVdS, BS, JL, LM, PM, JV, WD, ASD

Formal Analysis: RMJ, TVdS, BS, JL, DS, LMA

Investigation: RMJ, TVdS, BS, JL, RAS, DS, AVdL, EL, LMA, KVV, LM, NY, AW, RVT, RES, EZ, MWL

Resources: RES, EZ, MWL, MW, PM, JV, JO, MH, WD, ASD

Data curation: RMJ, TVdS, BS, JL, DS, LMA, JA

Writing – Original Draft: RMJ, TVdS, BS, WD, ASD

Writing – Review and Editing: RMJ, TVdS, BS, JL, RAS, DS, OD, AVdL, EL, LMA, RB, KVV, LM, NY, JA, AW, RVT, RES, EZ, MWL, MW, PM, JV, JO, MH, WD, ASD

Visualization: RMJ, TVdS, BS, JL, LMA, OD

Supervision: BS, PM, JV, JO, MH, WD, ASD

Project Administration: RMJ, TVdS, BS, WD, ASD

Funding Acquisition: TVdS, EL, PM, JV, JO, MH, WD, ASD

## Supplemental Figure Legends

**Figure 1 S1.**
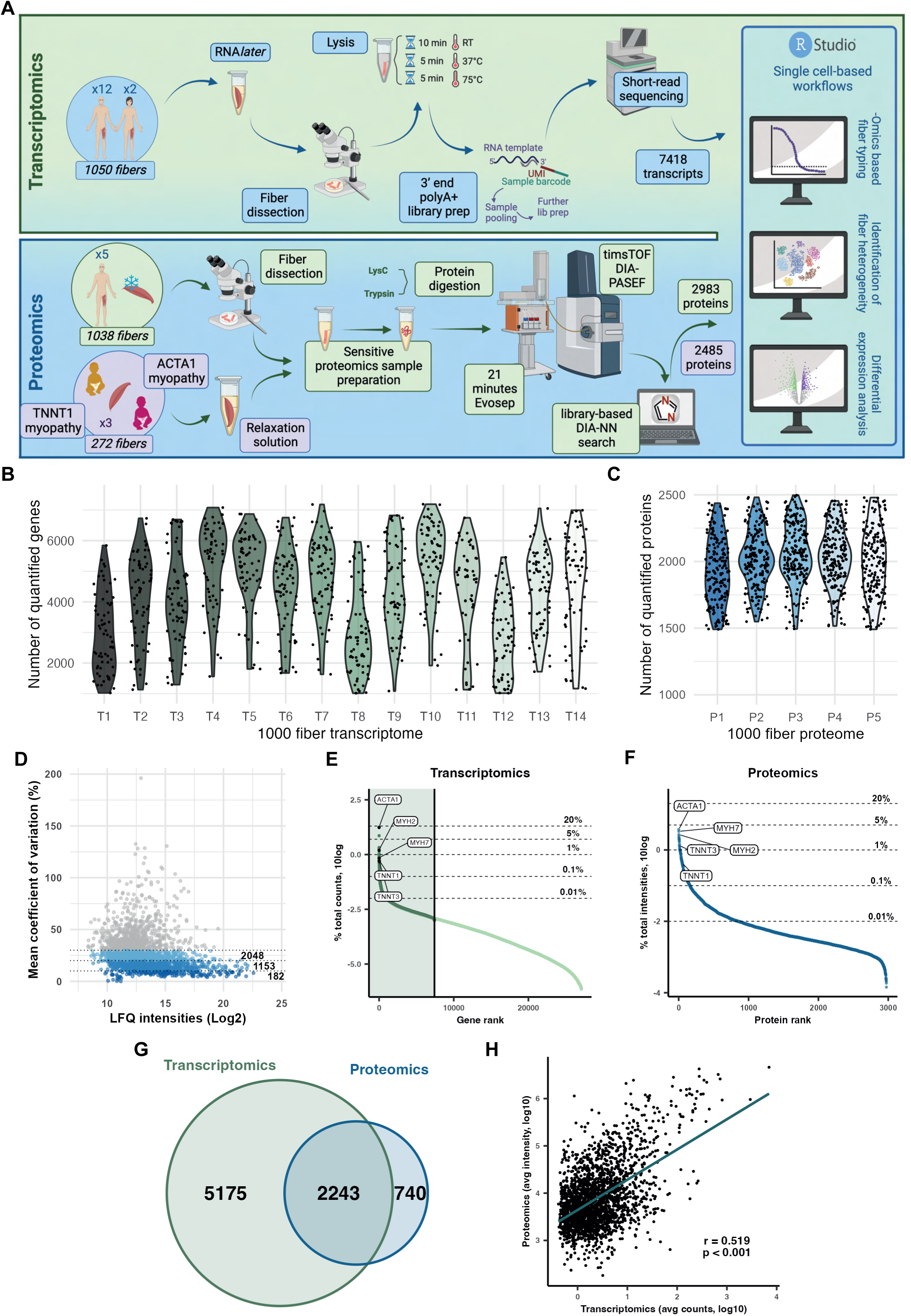
Quality control of novel transcriptomic and proteomic workflows. (**A**) Detailed schematic of developed workflows. (**B-C**) Number of quantified genes/proteins by participant and sample in 1000 fiber transcriptomics and proteomics. (**D**) Coefficient of variation by protein in 1000 fiber proteomics. (**E-F**) Dynamic range of genes in 1000 fiber transcriptomics and of proteins in 1000 fiber proteomics. (**G**) Overlapping features within 1000 fiber transcriptomics and proteomics. (**H**) Correlation between features within 1000 fiber transcriptomics and proteomics.

**Figure 1 S2.**
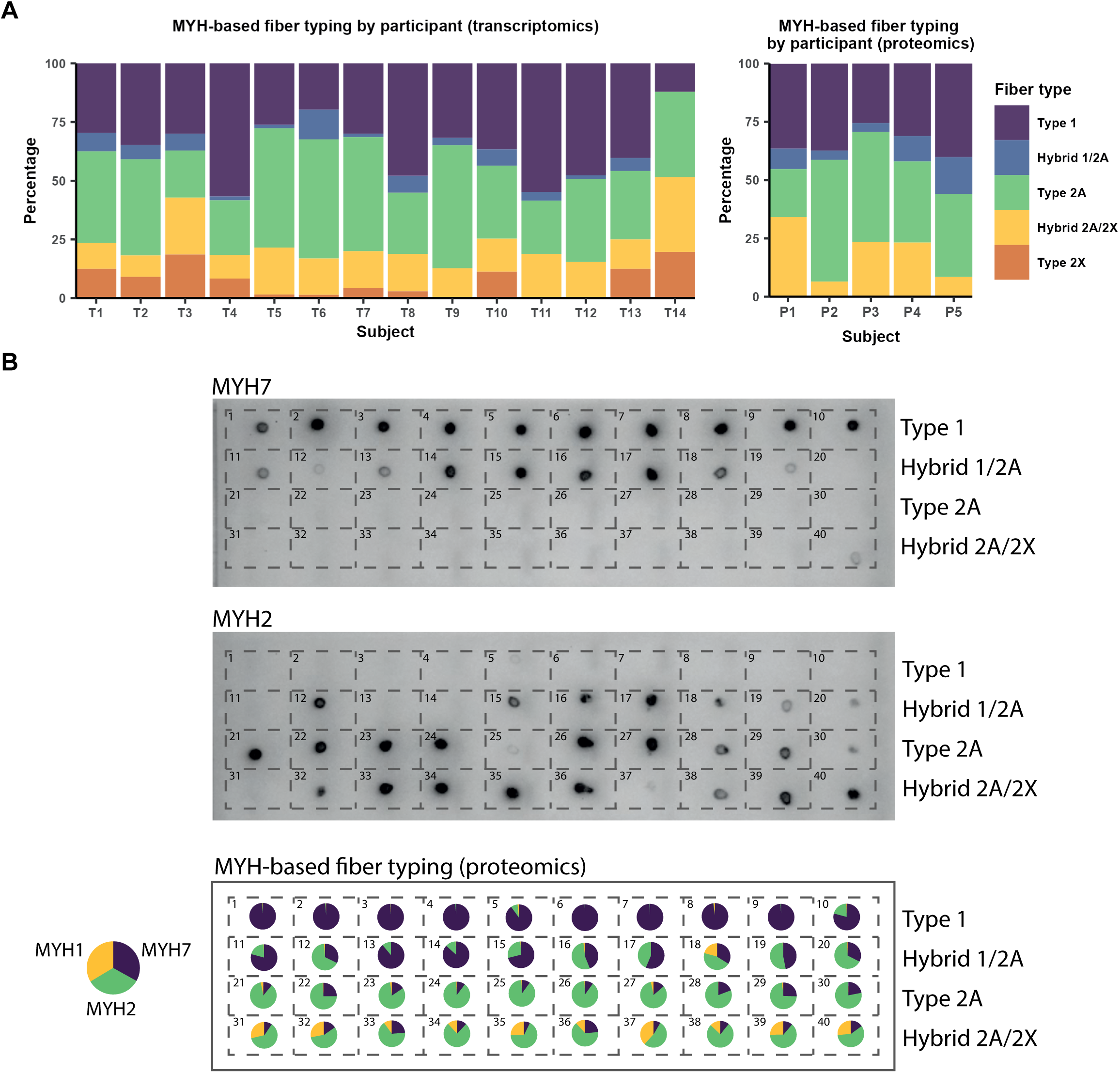
Higher sensitivity with proteomics compared to standard dot blotting. (**A**) MYH-based fiber type proportions by participant. (**B**) Comparison between dot blot- and proteomics-based fiber typing. The pie chart indicates the abundance for each MYH isoform in the single-fiber proteomics data.

**Figure 1 S3.**
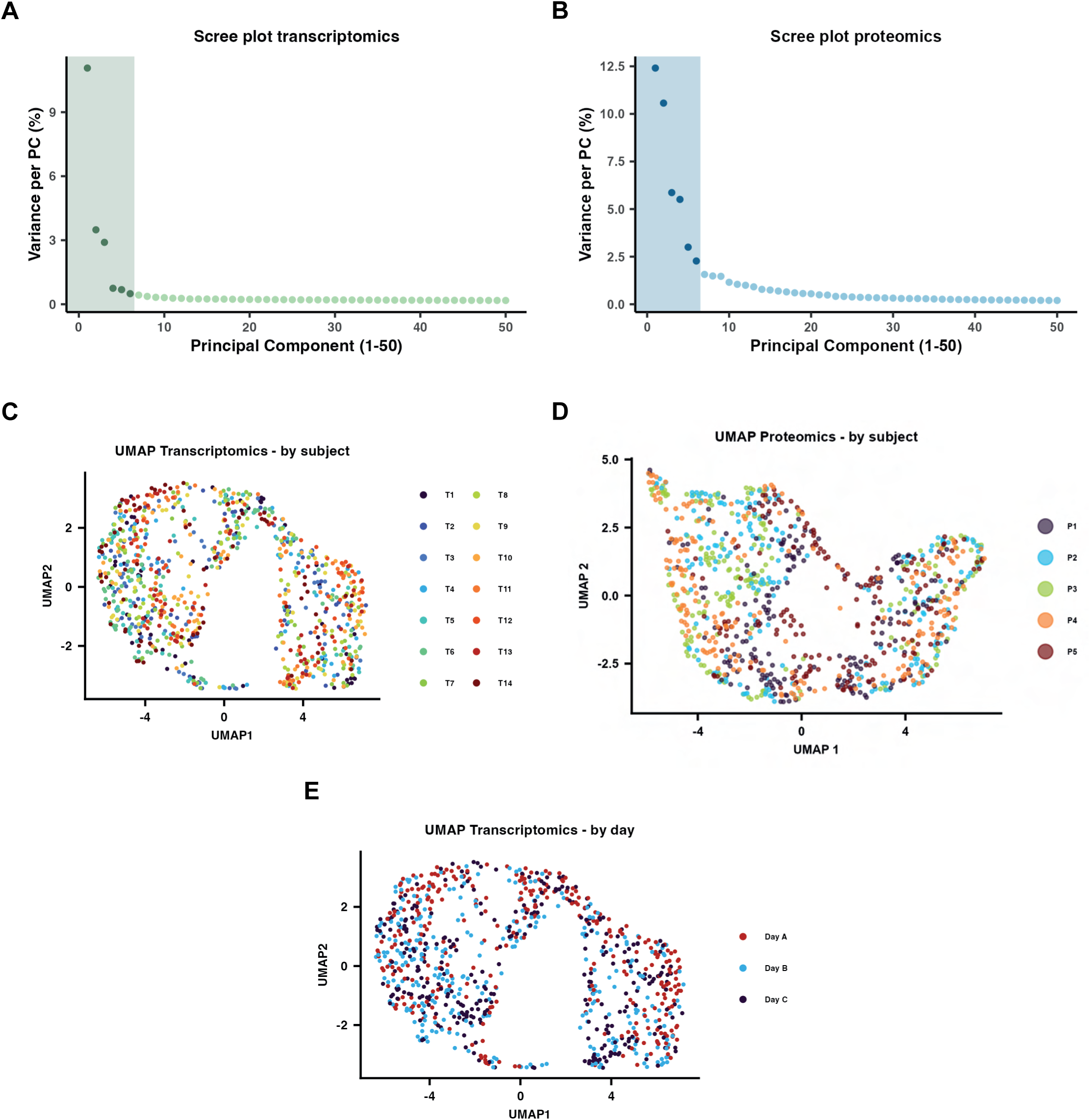
Clustering of transcriptomic and proteomic data. (**A-B**) Scree plots of transcriptomics and proteomics. Colored box indicates the number of principal components used for clustering. (**C-D**) 1000 fiber transcriptomics and proteomics UMAP colored by participant. (**E**) 1000 fiber transcriptomics UMAP colored by trial day.

**Figure 1 S4.**
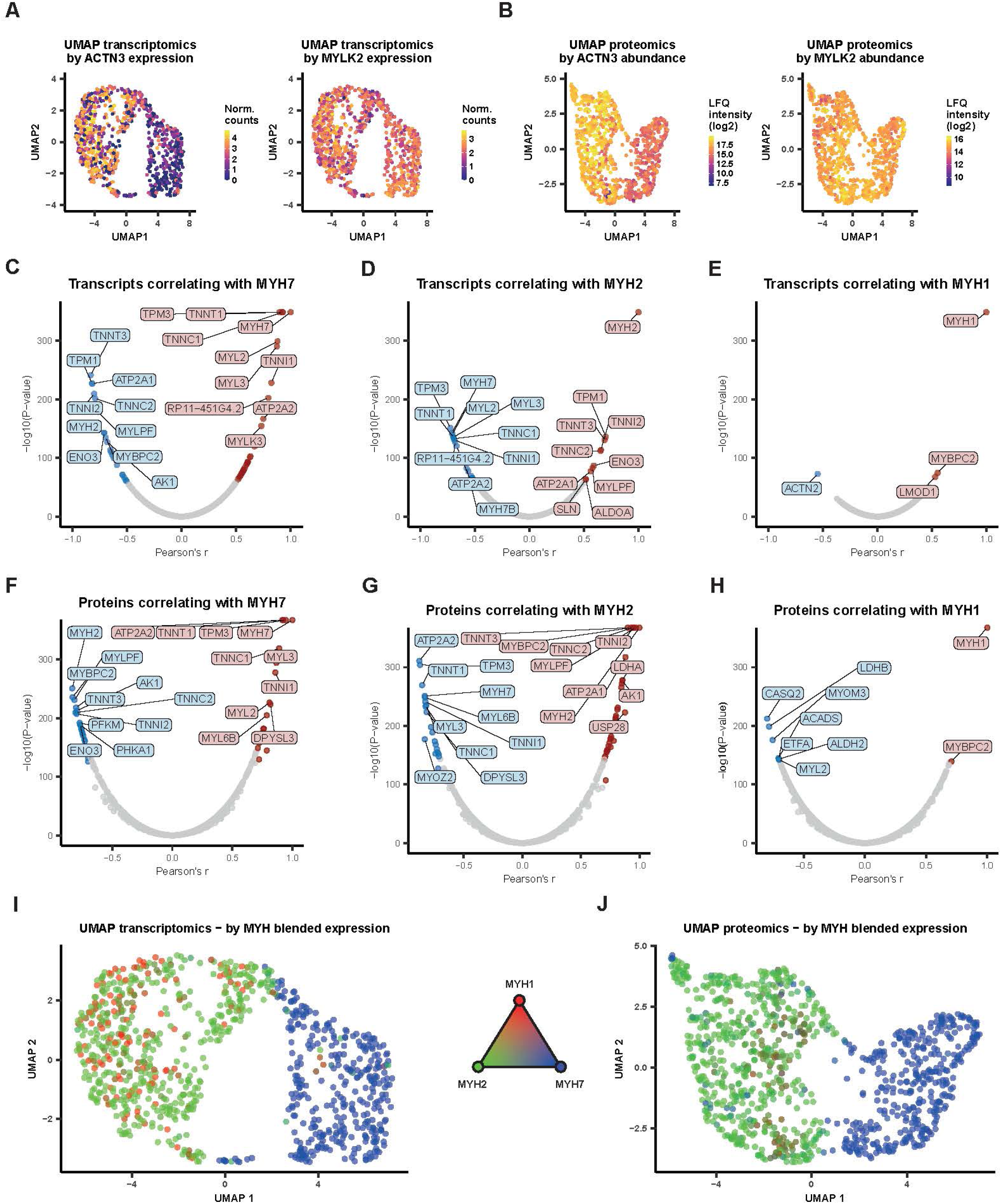
Type 2X is not a distinct fiber type. (**A-B**) Feature plots for ACTN3 and MYLK2 expression/abundance in transcriptomics and proteomics data. (**C-H**) U-plots demonstrating correlation of transcripts/proteins with the different MYH isoforms. X-axis depicts Pearson’s r coefficient, y-axis depicts -log10 of the p-value. (**I-J**) UMAP plots for transcriptomics and proteomics showing blended expression of MYH7 (blue), MYH2 (green) and MYH1 (red).

**Figure 2 S1.**
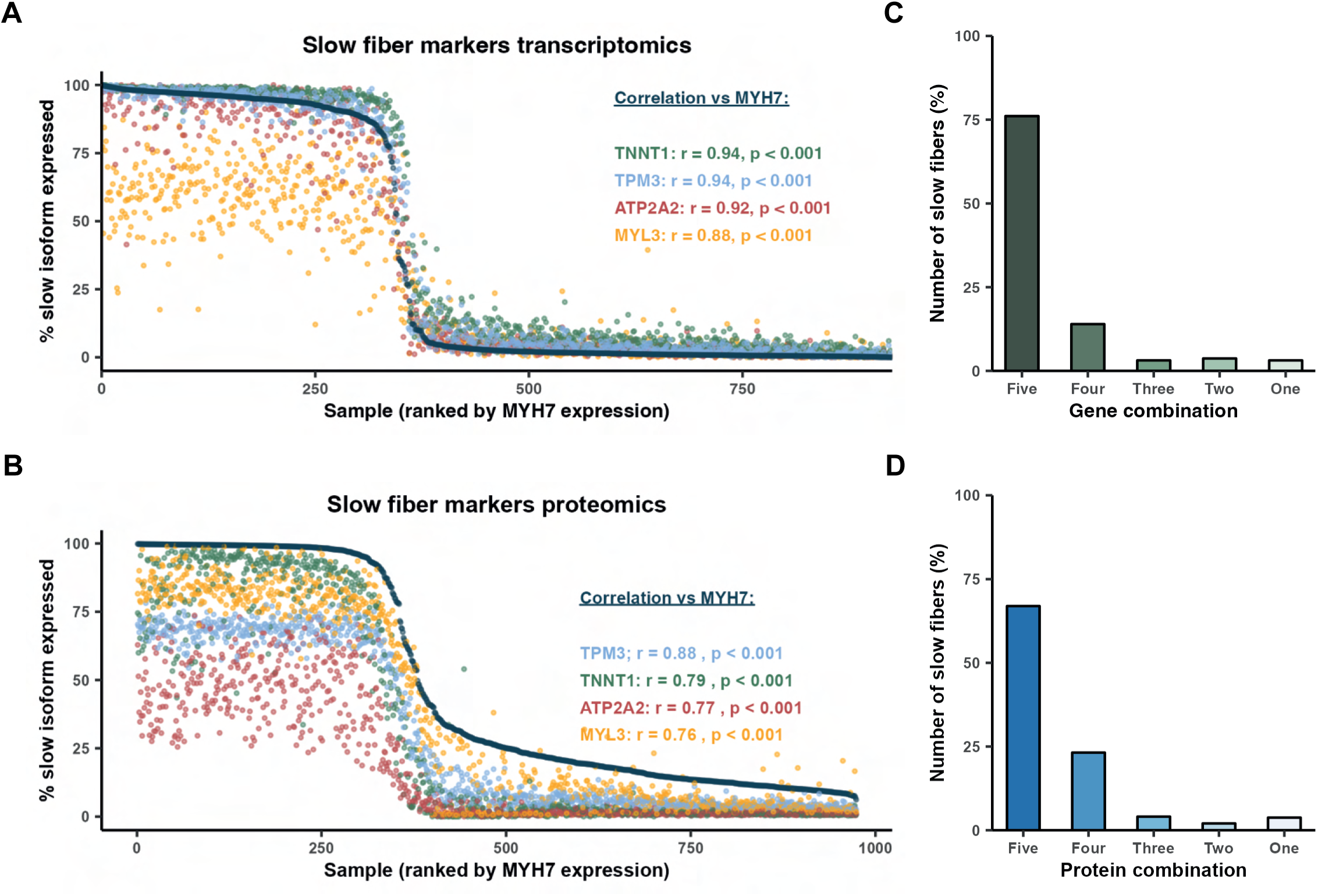
Heterogeneity of fibers with well-known fiber type specific features. (**A-B**) Fiber type curves of established slow fiber type genes/proteins in 1000 fiber transcriptomics and proteomics. (**C-D**) Proportion of fibers determined to be slow via different numerical combinations of genes/proteins (from A-B) in 1000 fiber transcriptomics and proteomics.

**Figure 2 S2.**
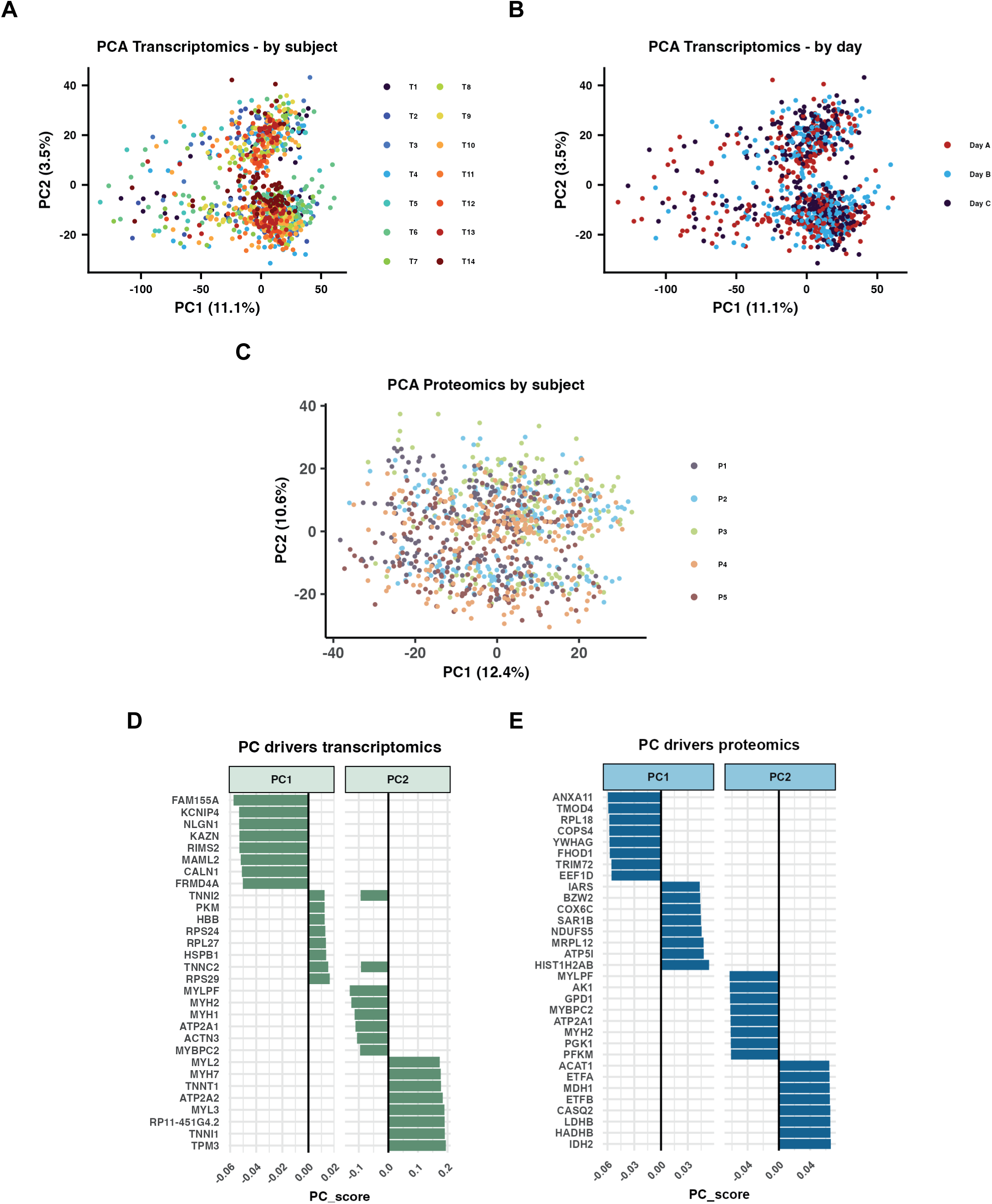
Additional data of Principal Component Analysis (PCA). (**A-B**) 1000 fiber transcriptomics PCA plot colored by participant or trial day. (**C**) 1000 fiber proteomics PCA plot colored by participant. (**D-E**) Top gene and protein drivers of PC1 and PC2.

**Figure 3 S1.**
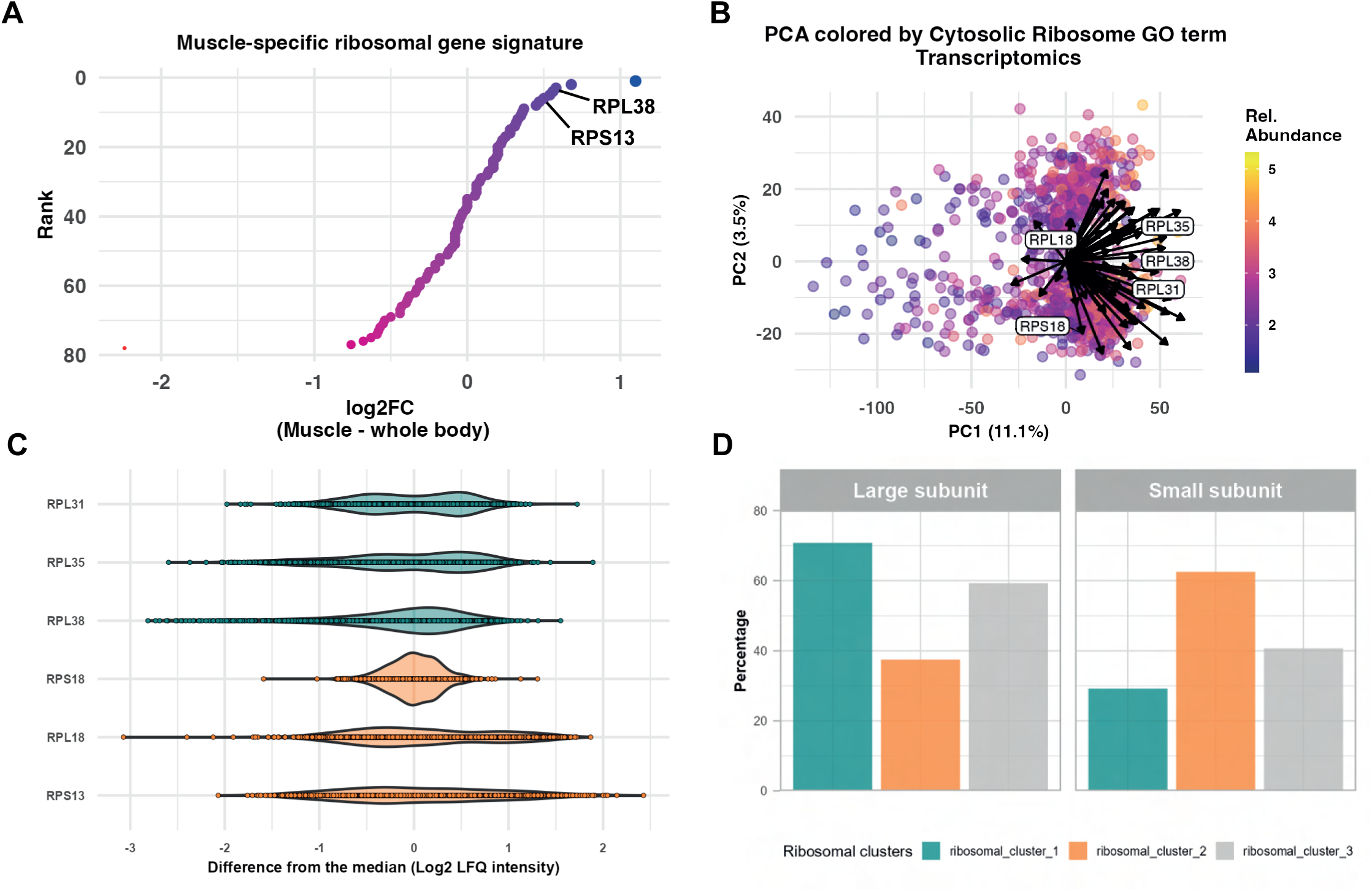
Ribosomal heterogeneity of skeletal muscle fibers. (**A**) mRNA expression of ribosomal proteins in skeletal muscle compared to the whole body. Data derived from Panda et al^67^. (**B**) PCA plot colored by cytosolic ribosome gene ontology (GO) term in the transcriptome. Arrows indicate the direction towards which the proteins are driving the variance in the PCA plot. The length of the lines represents the principal component score for a given protein. (**C**) Differences in protein abundance of ribosomal proteins in single muscle fibers compared to the median abundance. (**C**) Proportions of ribosomal proteins in each cluster by small (40S) and large (60S) subunits.

**Figure 4 S1.**
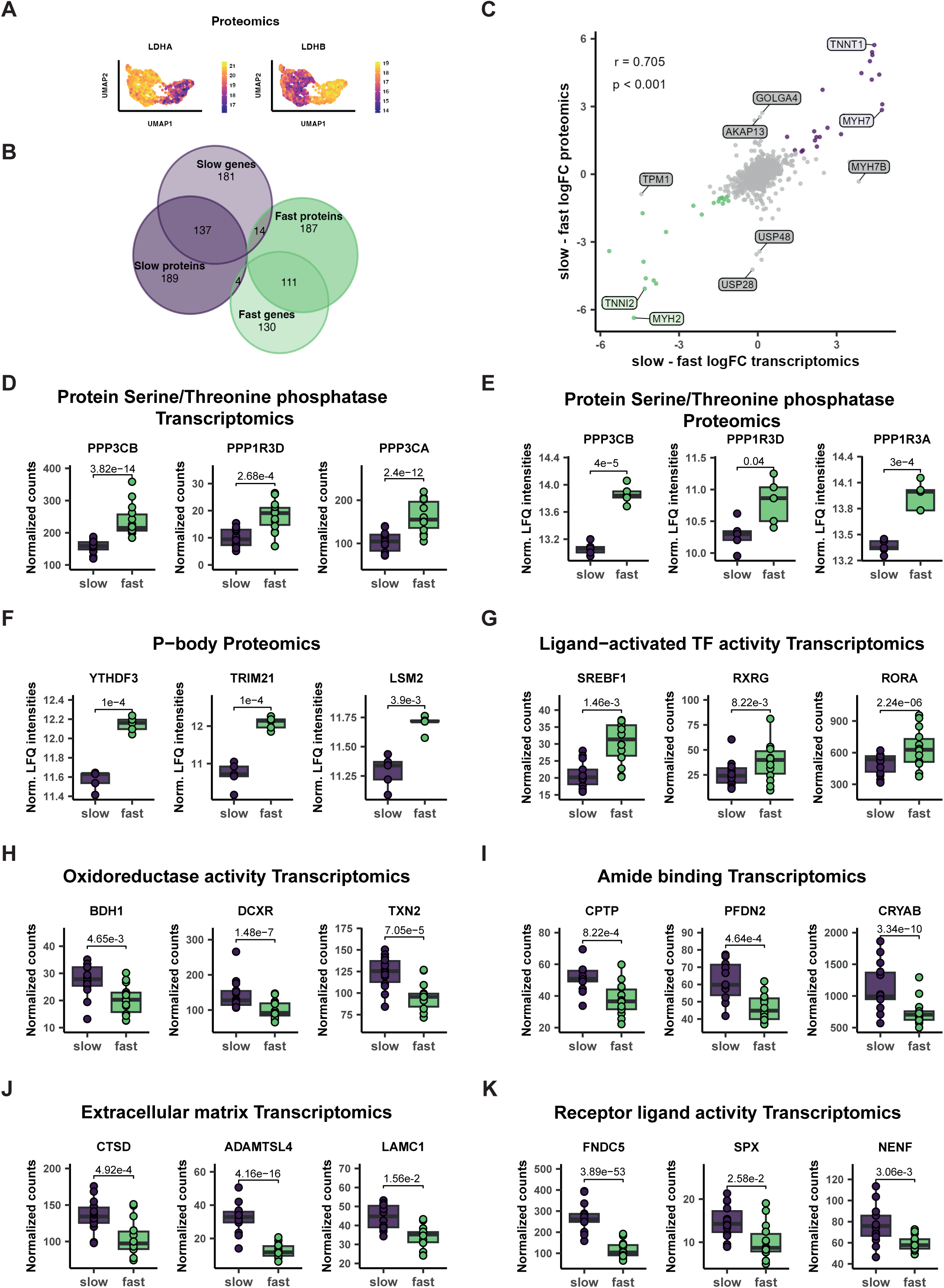
Gene and protein signatures of fast and slow fibers. (**A**) Feature plots of LDH protein isoforms between slow and fast fibers. (**B**) Overlap of significantly different genes/proteins between slow and fast skeletal muscle fibers in 1000 fiber transcriptomics and proteomics. (**C**) Correlation between the log_2_ fold change difference between slow and fast skeletal muscle fibers within 1000 fiber transcriptomics and proteomics. (**D-K**) Expression profiles of top drivers of selected enrichment terms within slow or fast fibers with indication of adjusted p-values.

**Figure 5 S1.**
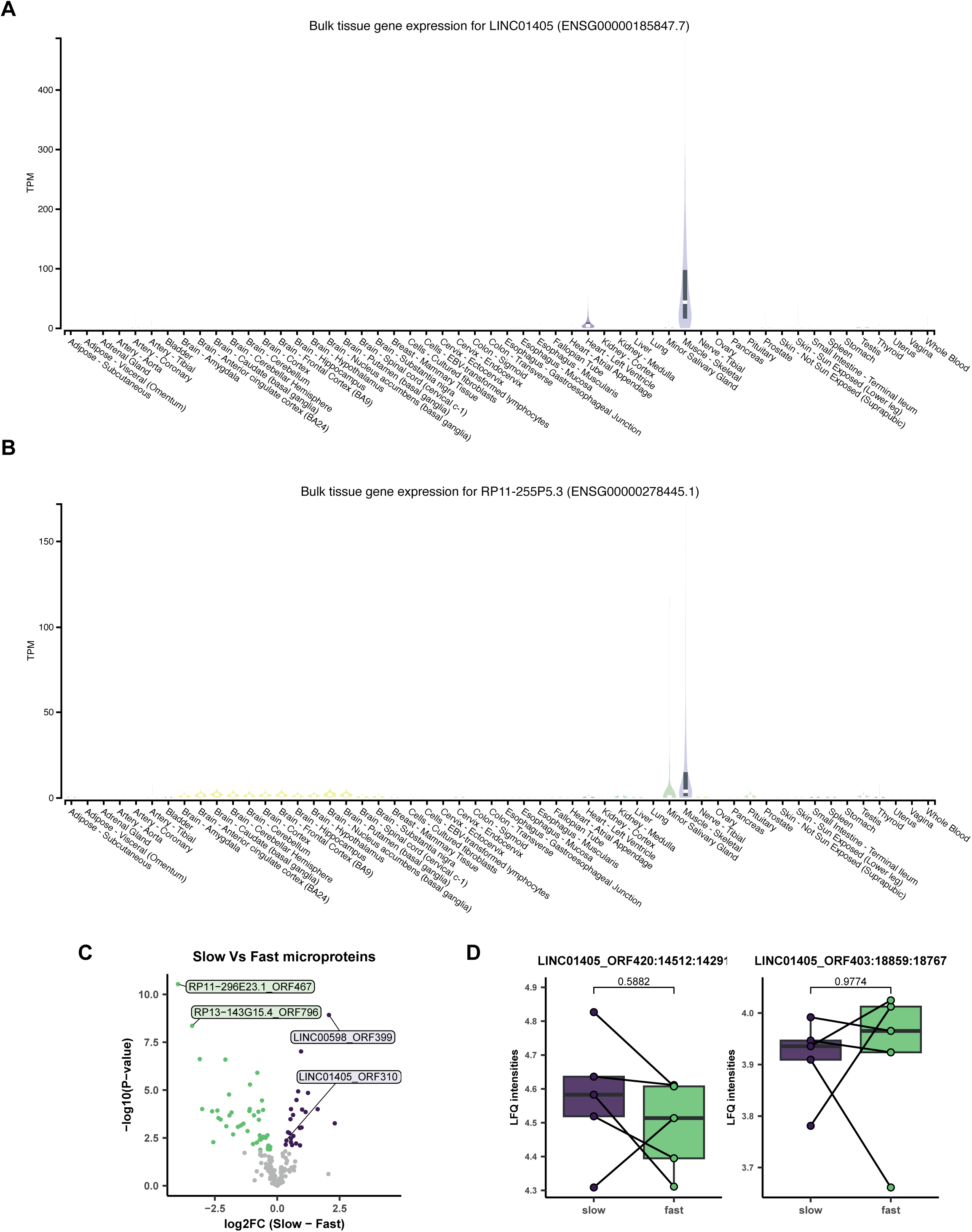
Specific expression of non-coding RNA and microproteins in skeletal muscle. (**A-B**) Expression profile of *LINC01405* and *RP11-255P5.3* in different tissues. (**C**) Volcano plots comparing microprotein abundance between the slow and fast clusters identified from the uniform manifold approximation and projection (UMAP) plots in Fig 1G-H. Colored dots represent significantly different microproteins at an FDR < 0.05. **(D)** Slow/fast fiber-type comparison of LINC01405_ORF52 14515:14294 and LINC01405_ORF503 18862:18770 microproteins.

**Figure 6 S1.**
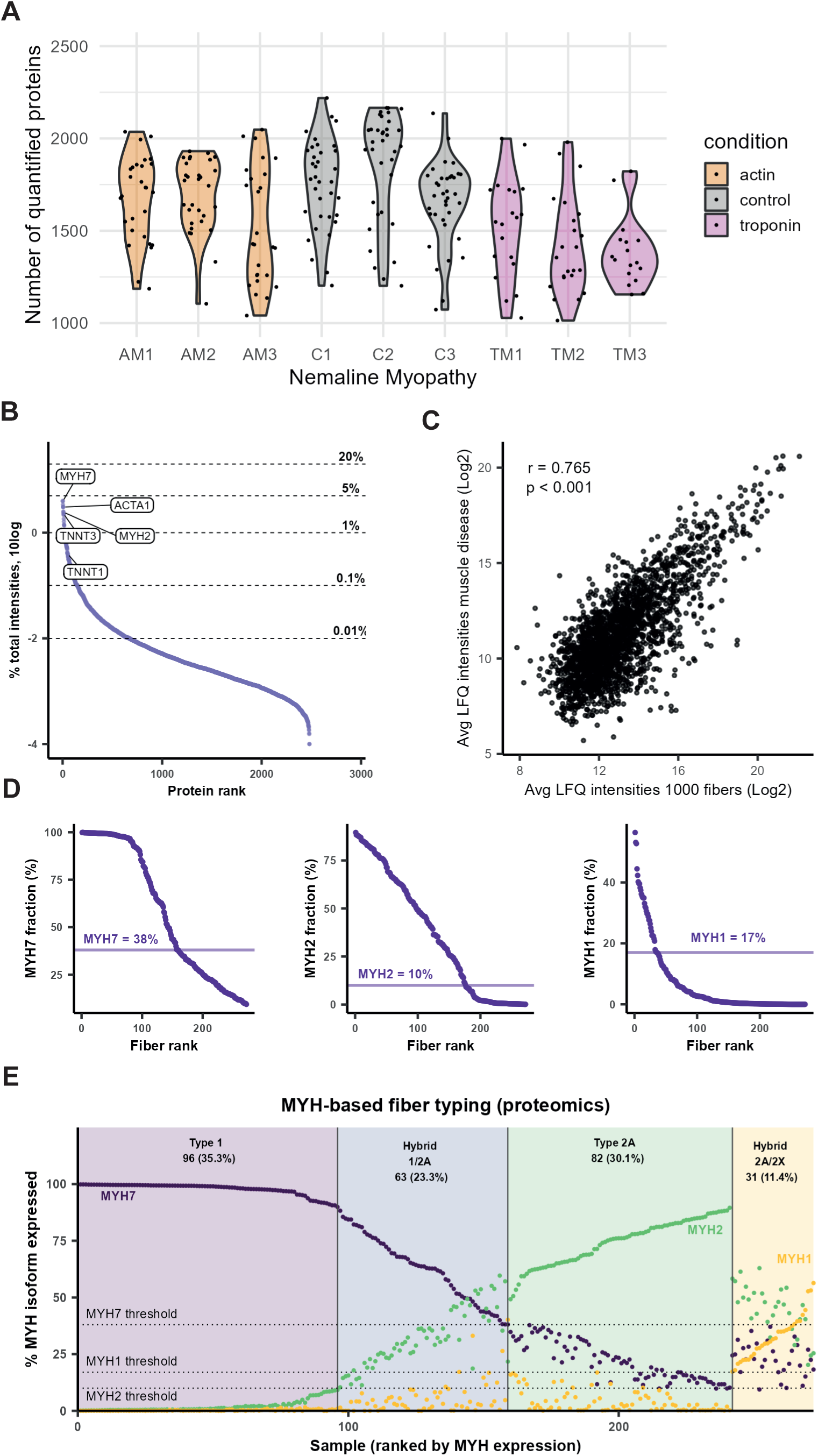
Quality control and fiber typing of nemaline myopathy dataset. (**A**) Number of quantified proteins by participant and sample in nemaline myopathy proteomics. (**B**) Dynamic range of proteins in nemaline myopathy proteomics. (**C**) Correlation between features within 1000 fiber proteomics and healthy controls of nemaline myopathy proteomics. (**D**) Dynamic range curves for MYH7, MYH2, and MYH1, with calculated assignment thresholds for fiber typing. (**E**) Distribution of fibers by expression of MYHs in nemaline myopathy proteomics.

**Figure 6 S2.**
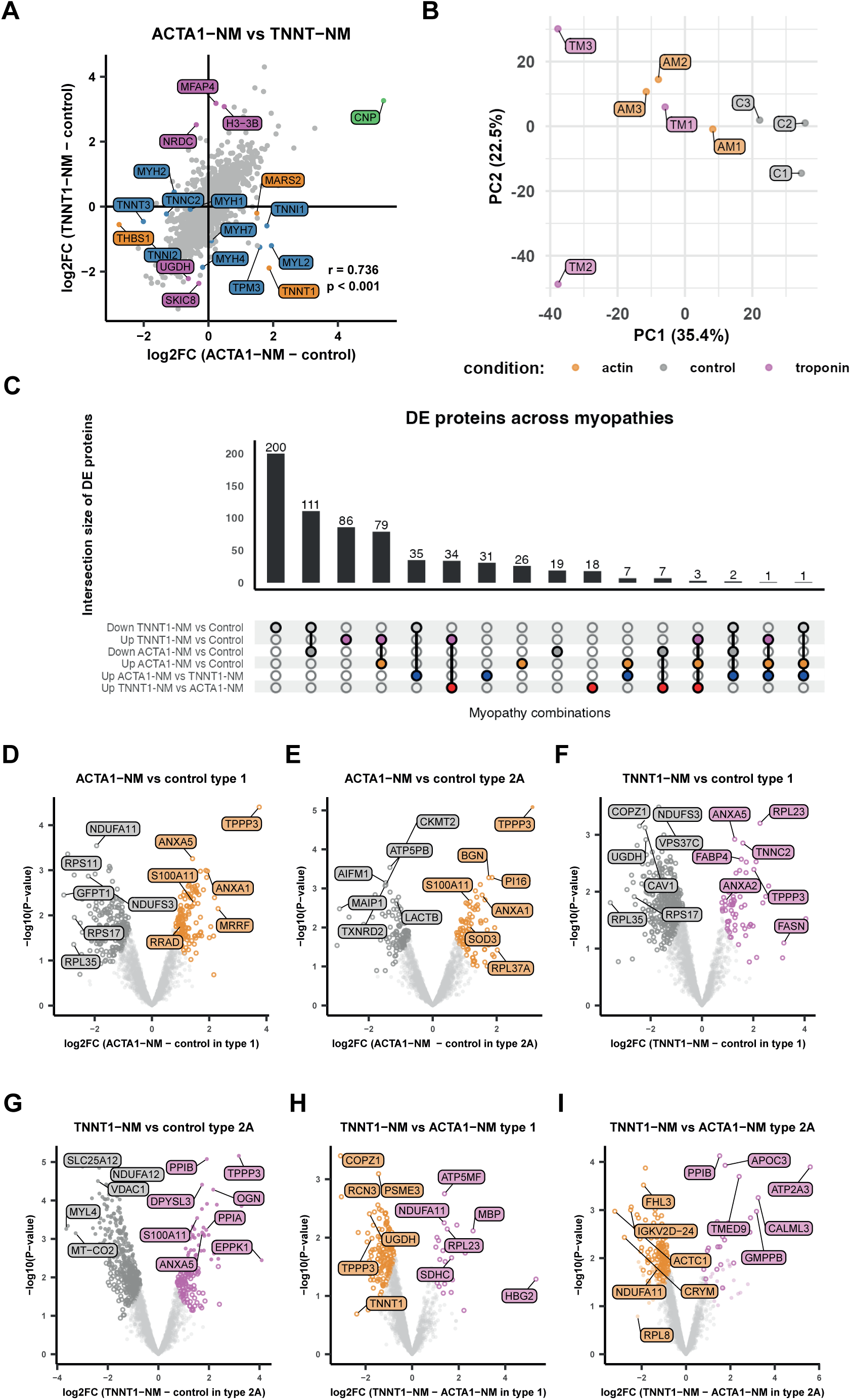
Fiber type-independent remodelling in nemaline myopathy. (**A**) Correlation between the log_2_ fold change difference between *ACTA1*- and *TNNT1*-nemaline myopathies with control participants. (**B**) PCA of nemaline myopathy patients (pseudobulked protein profile form single fibers). (**C**) Upset plot of significantly regulated proteins *ACTA1*- and *TNNT1*-nemaline myopathies. (**D-I**) Volcano plots comparing the *ACTA1*- and *TNNT1*-nemaline myopathies with control participants and between *ACTA1*- and *TNNT1*-nemaline myopathies by MYH-based fiber type. Colored circles represent significantly different proteins at π < 0.05, darker dots represent significantly different proteins at FDR < 0.05.

**Figure 6 S3.**
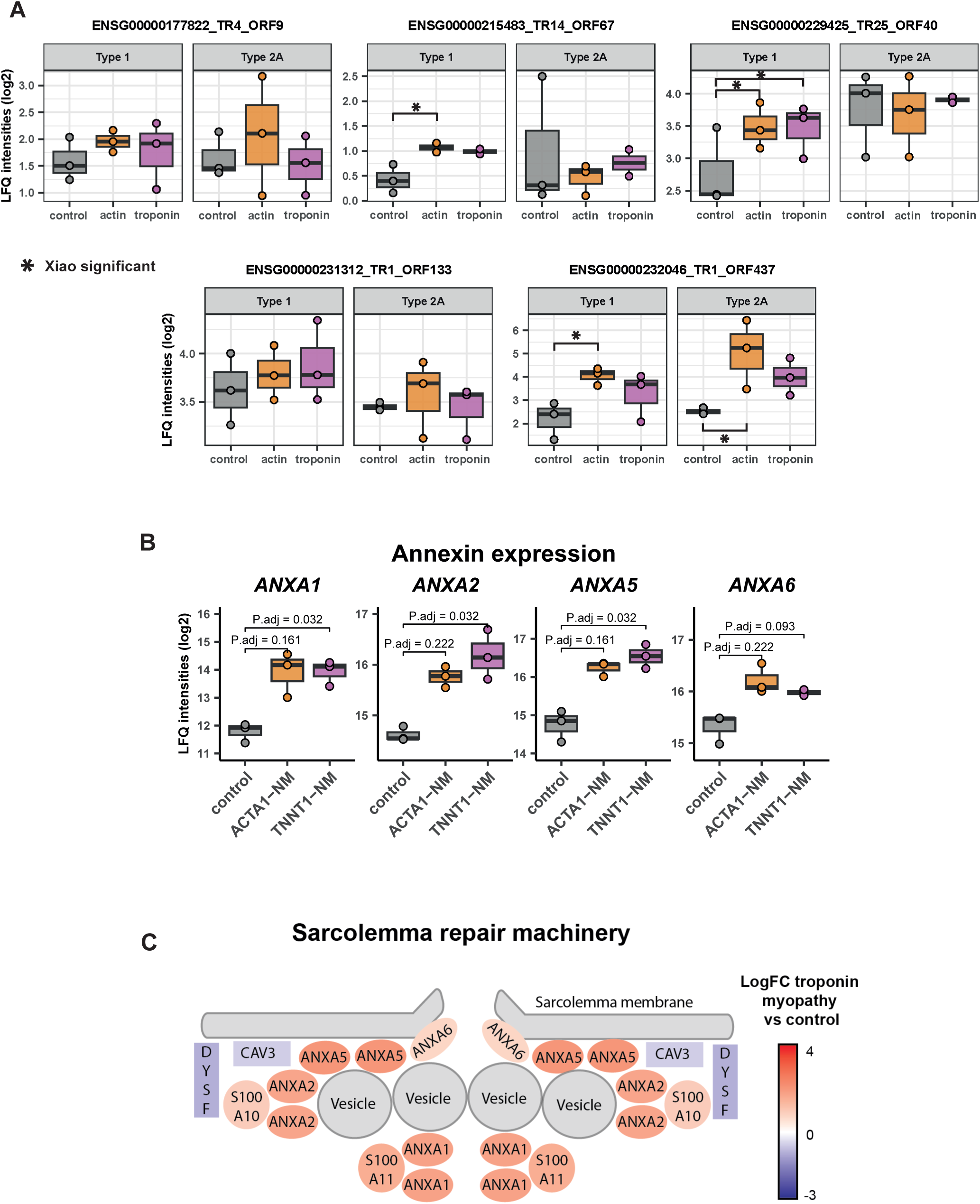
Effects of nemaline myopathy on microproteins and the sarcolemma repair machinery. **(A)** Protein abundance of identified microproteins within ACTA1-, TNNT1- and control participants. Significantly different comparisons between groups are highlighted. **(B)** Abundance of annexin proteins in *ACTA1*- and *TNNT1*-nemaline myopathy. (**C**) Schematic of changes within the sarcolemma repair machinery in nemaline myopathy.

**Figure 6 S4.**
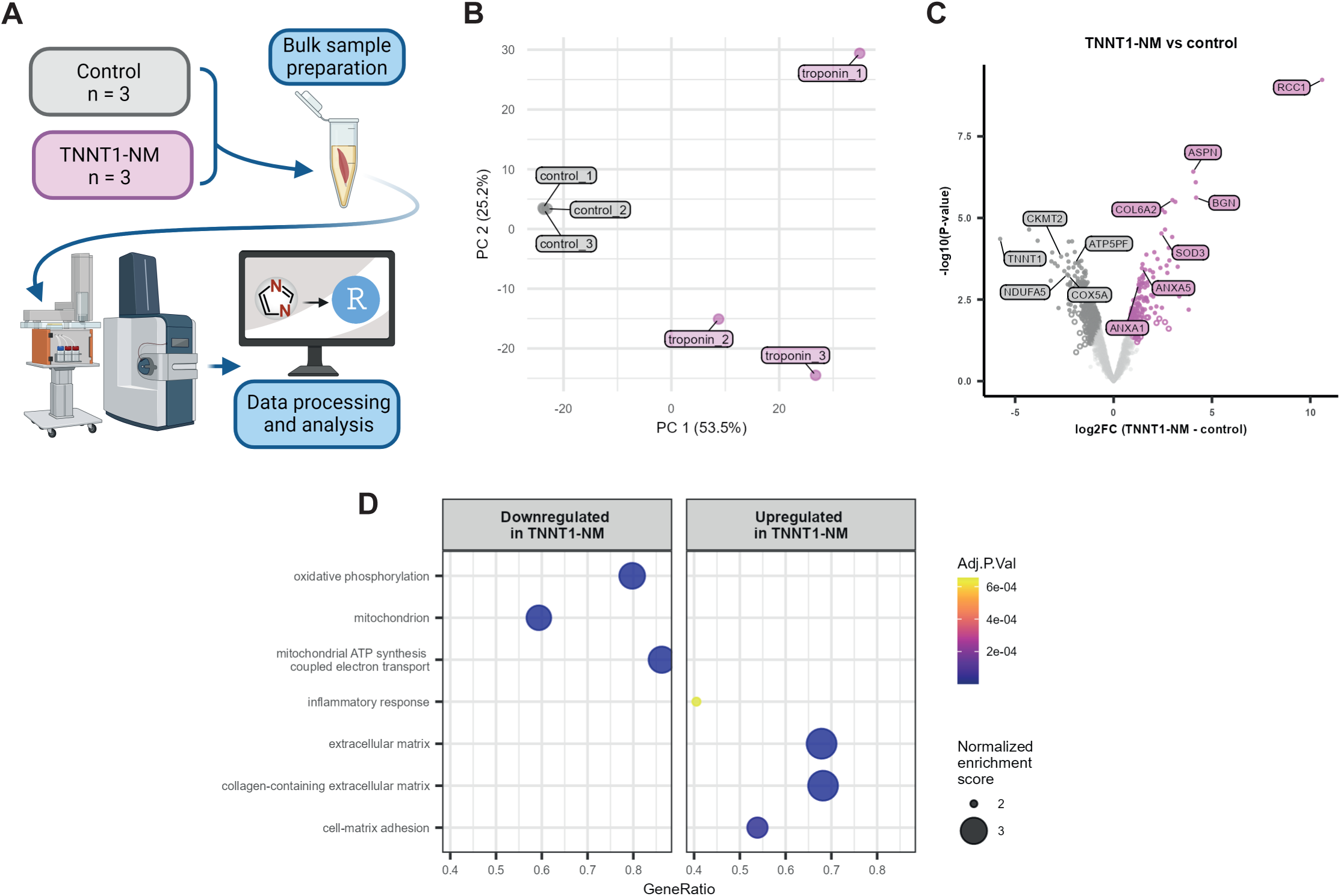
Bulk muscle proteomics recapitulates single-fiber analysis of nemaline myopathy samples. (**A**) Workflow of bulk muscle proteomics for control participants and *TNNT1*-nemaline myopathy patients. (**B**) Principal component analysis (PCA) plot of bulk skeletal muscle from *TNNT1*-nemaline myopathy patients and controls. **(C)** Volcano plot comparing the *TNNT1*-nemaline myopathy with control participants. Colored circles represent significantly different proteins at π < 0.05, darker dots represent significantly different proteins at FDR < 0.05. **(D)** Enrichment analysis of significantly different proteins in the bulk muscle proteome between *TNNT1*-nemaline myopathy patients and control participants.

